# Friend Request Accepted: Fundamental Features of Social Environments Determine Rate of Social Affiliation

**DOI:** 10.1101/2025.02.05.636557

**Authors:** Sankalp Garud, Miruna Rascu, Sorcha Hamilton, Ingrid Yu, Matthew F.S. Rushworth, Miriam C. Klein-Flügge

**Affiliations:** Wellcome Centre for Integrative Neuroimaging, Department of Experimental Psychology, University of Oxford, Oxford OX1 3SR, United Kingdom; Department of Psychiatry, Warneford Hospital, University of Oxford, Oxford OX3 7JX, United Kingdom; Department of Psychology, University of Bath, Bath BA2 7AY, United Kingdom

## Abstract

Humans start new friendships and social connections throughout their lives and such relationships foster mental and physical well-being. While friendship initiation may depend on alignment of subtle and complex personal variables, here we investigated whether it also depends on basic features of social environments. This would be analogous to other fundamental behaviours like foraging which depend on basic features of the environment such as the density of opportunities and the likelihood of success. In a pre-registered online study (n=783), we found people were more likely to send friend requests as the density of friendship opportunities decreased and frequency of success increased. Further, we found task-related measures, like overall friend requests, were correlated with personality-related factors like social thriving and anhedonia. Next, in an ultra-high-field fMRI study (n=24), we found that both fundamental features of social environments – opportunity density and frequency of success – affected neural activity across a network of regions linked to foraging including dorsal raphe nucleus, substantia nigra, and anterior insula. Finally, in resting-state fMRI data (n=400), we showed that model predicted estimates of anhedonia were related to functional connectivity between components of the same network. Thus, humans consider the background statistics of an environment while making social decisions and these decisions are linked to activity in ancient subcortical circuits mediating the influence of environmental statistics on other aspects of behaviour. Moreover, individual differences in how environmental features influence social behaviour are associated with variation in personality and psychiatric traits, offering new insights into inter-individual variability in social functioning.

## INTRODUCTION

No one is a stranger to the warm feeling of connecting with people they love. Indeed, there is empirical evidence for a positive link between social connection and well-being. For instance, increased social contact is a major predictor of happiness, whereas increased loneliness and isolation are linked to feelings of depression and increased mortality risk^1–3^. Given the influence of social connection on our well-being, it is important to understand how these social connections start and what neural mechanisms support their formation.

Social psychologists have shown that similarity, familiarity, and proximity facilitate social connection between individuals^4^. It has also been argued that social connection might be a basic human need and even that, at the level of the brain’s motivation circuits, the impact of social deprivation resembles that of food deprivation^5,6^. If social connection is as much of a “fundamental need” as is food, then insights about how we initiate social contact might be gleaned from how humans and animals forage for food. For example, classical and modern studies of foraging have suggested that beyond the value of a food opportunity in itself, animals take into account background statistics of opportunities in an environment (like average rate of reward) when deciding whether to engage with any opportunity^7–9^. In an analogous manner, we hypothesised that decisions about whether to initiate social engagement with an individual might depend on similar fundamental features of the background statistics of social environments.

In the present study, we investigated human decisions to initiate affiliation with other individuals in different environments. Drawing on investigations into foraging^7–9^, we hypothesised that two features of social environments would influence people’s decisions to initiate friendships: (i) the average friendliness of an environment, that is, the average success of one’s attempts to create friendships in a given environment, and (ii) the opportunity density, that is, the number of opportunities one gets to create friendships in a given environment.

These predictions stem from findings that animals choose to allocate more resources in energy rich environments and tend to be more selective in denser patches^10^. Therefore, we predicted that, by analogy, people would attempt to initiate more friendships in friendlier environments as compared to hostile environments (i.e. they would expend more resources pursuing opportunities when each opportunity is, in itself, likely to be richer, for example, in environments that are richer on average), and initiate fewer friendships in denser environments as compared to sparser environments (i.e. they become more selective in denser patches). If people were disinclined to initiate friendships in dense or hostile environments, then, even on occasions when they do initiate a friendship attempt, their reaction times (RTs) might still be slower. By contrast, RTs when participants avoid initiating friendships might be longer in sparse and friendly environments (all behavioural hypotheses were pre-registered at https://osf.io/62hw7).

Given the link between social isolation and mental well-being^1–3,11^, we also carried out a second line of investigation, examining the relationship between the impact of the statistics of social environments on people’s tendencies to initiate social contact and their mental health and personality profile. To assess state and trait mental health, we asked people to complete more than 100 items from multiple standardised questionnaires that measure symptoms for conditions ranging from depression to anxiety and loneliness. We then used factor analysis to define mental health dimensions from correlated subscales corresponding to traditional mental health categories^12^, and investigated the relationship between these transdiagnostic factors and the impact that the statistics of social environments had on seeking social affiliation.

A third line of investigation considered the neural mechanisms through which affiliation decisions were influenced by statistics of the social environment. We focused on a cortico-subcortical circuit implicated in the tracking of the background statistics of reward environments during foraging^8,9,13–19^ to test whether they played an analogous role in social affiliation seeking. It has been argued that opportunity costs associated with the average rate of gustatory rewards in rats are tracked by tonic dopamine-linked midbrain nuclei such as the substantia nigra (SN)^20^. Relatedly, it has been suggested that neural activity spanning a distributed network comprising anterior insula (aI) and habenula (Hb) reflects the evaluation that a monetary opportunity has on behaviour, given both its own value and the background context, and that this evaluation is then translated into a decision to act via SN^16^. We reasoned that the same circuit might mediate the influence of the social environment on whether to engage with any particular social opportunity. Other studies have emphasized another neuromodulatory system centred on the dorsal raphe nucleus (DRN), in tracking the average value of the environment^18,21^ or transitions in its average value^19^ . As is the case for SN, DRN activity patterns also emerge in the context of interactions with aI and Hb^17–19,21^. In addition, given its role in tracking and individuating social agents^22–30^, we also examined activity in dorsomedial prefrontal cortex (dmPFC). We focused on a predefined area of interest in dmPFC in area 9^31^.

In summary, we hypothesised that decisions to initiate social contact might be influenced not just by the opportunity itself but also by statistics of the social context. If this is the case, then it would suggest social contact-seeking might operate in an analogous manner to the seeking of food or monetary rewards in humans and other animals. We, therefore, at a neural level, investigated activity across a distributed cortico-subcortical circuit comprising DRN, SN, Hb, dmPFC and aI that has been linked to the tracking of reward statistics and the influence they exert over behaviour. We also examined how variation in social affiliation seeking might be linked to variation in personality and psychiatric traits.

## RESULTS

### Environmental friendliness and density influence social affiliation choices

783 participants performed a Friend Request Task (Fig.1) in which they were asked to imagine moving to a new city and making new friends. They were told they would be helped to create connections by being taken to different social clubs which varied in two respects: the friendliness of the people in the club and the number of people that appeared in the club. As a result, four combinations of clubs were possible: friendly-densely populated (FD), friendly-sparsely populated (FS), hostile-densely populated (HD), and hostile-sparsely populated (HS) (Fig.1b). Participants spent 2.5 minutes, which we refer to as a block, in one club before moving on to another. In a friendly club, 80% of all friend requests that a participant sent were accepted whereas in a less friendly (more hostile) club, only 20% of all requests were accepted. In a densely populated club, the time between two consecutive encounters with club members (being shown a club member face) was briefer; the faces appeared every 2 seconds with a jitter between 400 and 700 milliseconds. In a sparse club, the time between consecutive faces was 5 seconds with an equivalent jitter. The background colour associated with each club (blue or green counterbalanced over participants) indicated the friendliness level of the club and the background pattern (smaller or larger circles counterbalanced over participants) indicated the density of club encounters. Inside each club, participants were shown a series of faces (each face was shown only once) and given the opportunity to send or skip sending a friend requests by pressing one button or another. If participants sent a friend request, the next screen would show them whether their request was accepted or rejected (Fig.1a). There was no consequence (monetary or temporal) associated with either action (skipping or requesting) or outcome (accept or reject). Twelve blocks in total were presented in a randomised order (three per block type).

**Figure 1.**
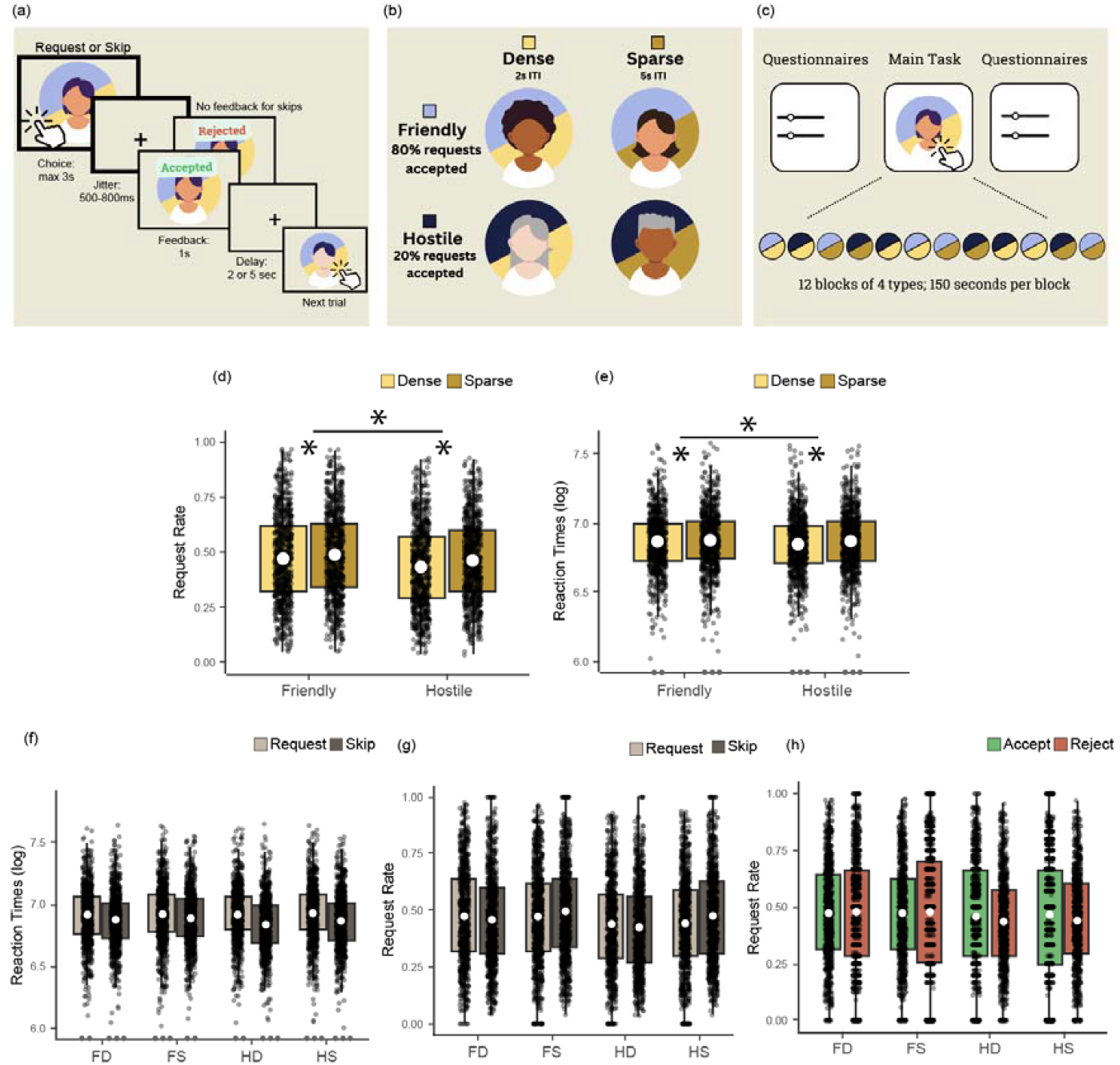
Task structure and behavioural effects in the Friend Request Task. (a) schematic representation and timings of a trial. (b) Two experimental manipulations, friendliness and density, were used. In a friendly block, 80% of friend requests sent are accepted, while in a hostile block 20% of requests are accepted. In a dense block, the inter-trial interval is a shorter 2s compared to the 5s interval in a sparse block. (c) Overview of the experimental structure in any given run. (d) Boxplots indicate the effects of friendliness and density on request rates (white circle indicates mean; box boundaries indicate interquartile range [IQR], encompassing the middle 50% of the data; whiskers extend to furthest data points within 1.5 × IQR from the box boundaries). The x-axis indicates the two levels of friendliness, and y-axis indicates the request rate (average number of requests sent) for an individual in that block type. Colour indicates the two levels of density. This shows higher request rates in sparser and friendlier environments. (e) The effects of friendliness and density on RTs. Y-axis indicates average log-transformed RTs in the respective block type. RTs are slower in friendly and sparse blocks. Dots shown at the y-axis limit denote outliers. (f) RTs split into request and skip trials. Colour indicates action (request/skip) (g) The effect of previous trial action on choice in the next trial. Colour indicates action (request/skip). (h) The effect of previous trial feedback on choice in the subsequent trial in different environment types. Colour indicates feedback received (accept/reject)

We first examined if the basic features of the social environment manipulated in our task affected people’s tendency to send friend requests. A 2x2 ANOVA (two levels of friendliness and two levels of density) revealed a significant effect of friendliness on request rates (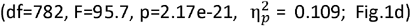). People were more likely to send friend requests when they were in a friendly environment as opposed to a hostile environment. Similarly, density also had a significant effect on request rates (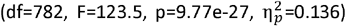). As such, people were more likely to send friend requests when they were in a sparse environment compared to a dense environment. Finally, the interaction between friendliness and density was significant (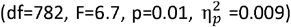). People were more likely to send friend requests in sparse environments, but particularly so when in hostile blocks. In a post-hoc analysis, we tested the robustness of these effects by excluding the first 10 trials from every block to allow for a brief transition period in which participants could experience the density of the new social environment they had just entered. We found that both friendliness and density effects remained significant (friendliness: 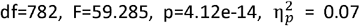; density: 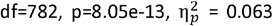; interaction: 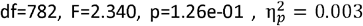;). Pre-registered mixed models showed the same results (see supplementary information).

Having characterized choice preferences in different social contexts, we next asked whether RTs similarly reflected features of the social environment. An analogous 2x2 ANOVA with factors friendliness and density identified a significant effect of friendliness on RTs (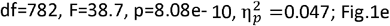); people were slower to respond in friendly compared to hostile environments. There was also a significant effect of density (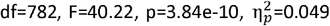); people were slower to respond in sparse compared to dense environments. There was an interaction between friendliness and density (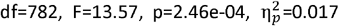). People were slower in sparse compared to dense environments, and this effect was especially pronounced in hostile blocks.

Next, we examined whether the action (request or skip) participants had chosen on the previous trial affected participant behaviour on the next trial and how this differed between social environments. Participants’ action on the current trial indeed affected RTs on the next trial (2 x 2 x 2 ANOVA: main effect of action: 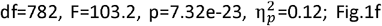). People were faster if the response was a skip as opposed to a request. There was also an interaction between the response type and friendliness (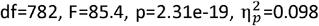). People were faster to skip than to request, but this effect was stronger in hostile blocks compared to friendly blocks. Similarly, there was an interaction between response type and density (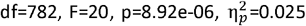). People were faster to skip than to request, but this effect was stronger in dense compared to sparse blocks. Thus, RTs also reflected key features of the social environments manipulated in our task.

The previous action (request or skip) also affected how likely people were to send friendship requests on the next trial. An analogous 2 x 2 x 2 ANOVA on request rates (friendliness x density x previous choice) showed a significant interaction between previous choice and density (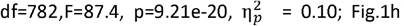). People were more likely to send a request following a previous request in dense environments, but they were more likely to send a request following a skip in sparse environments. The equivalent test for friendliness interacting with previous action was not performed as the result was not significant in our exploratory analysis, and therefore not pre-registered.

Finally, we examined whether social feedback received in one trial differentially affected how likely people were to send friendship requests in the following trial in the different social contexts. We extended the 2x2 ANOVA with friendliness and density to include a third factor of previous trial feedback (“accept” vs “reject”). Indeed, there was a main effect of previous trial outcome showing that the outcome received on a previous trial also affected participant behaviour in the next trial (2 x 2 x 2 ANOVA with factors friendliness, density, and previous trial feedback: previous outcome:(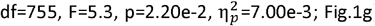). People were more likely to send a request after their previous request was accepted compared to when it was rejected. The interaction between friendliness and previous trial feedback was also significant (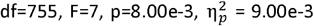); people were more likely to send a request following an acceptance, but this was particularly the case in hostile blocks. The equivalent test for feedback interacting with density was not performed because the result was not significant in our exploratory analyses performed in the preregistration cohort, and therefore not pre-registered.

### Transdiagnostic personality factors are associated with social connection seeking on the Friend Request Task

In addition to the behavioural task, the same 783 participants completed a series of standardised questionnaires to assess their personality and psychiatric profile (Methods). After analysing an initial exploratory dataset (N=206), we pre-registered our hypothesis (https://osf.io/jf6vs/) that it would be possible to identify seven social and non-social factors including indices of Social Thriving and Sensation Seeking that would be related to social connection seeking in the Friend Request Task. Six out of the seven pre-registered factors were observed in the confirmatory dataset: Social Thriving, Obsession/Compulsion, Impulsivity, Pleasure (reduced anhedonia), Sensation Seeking/Urgency, and Depression-Anxiety (Fig.2). We did not, however, identify a factor resembling the Social Pain Factor that we had found in the discovery data set analysis although, instead, we found some evidence for a factor that we termed social assurance.

**Figure 2.**
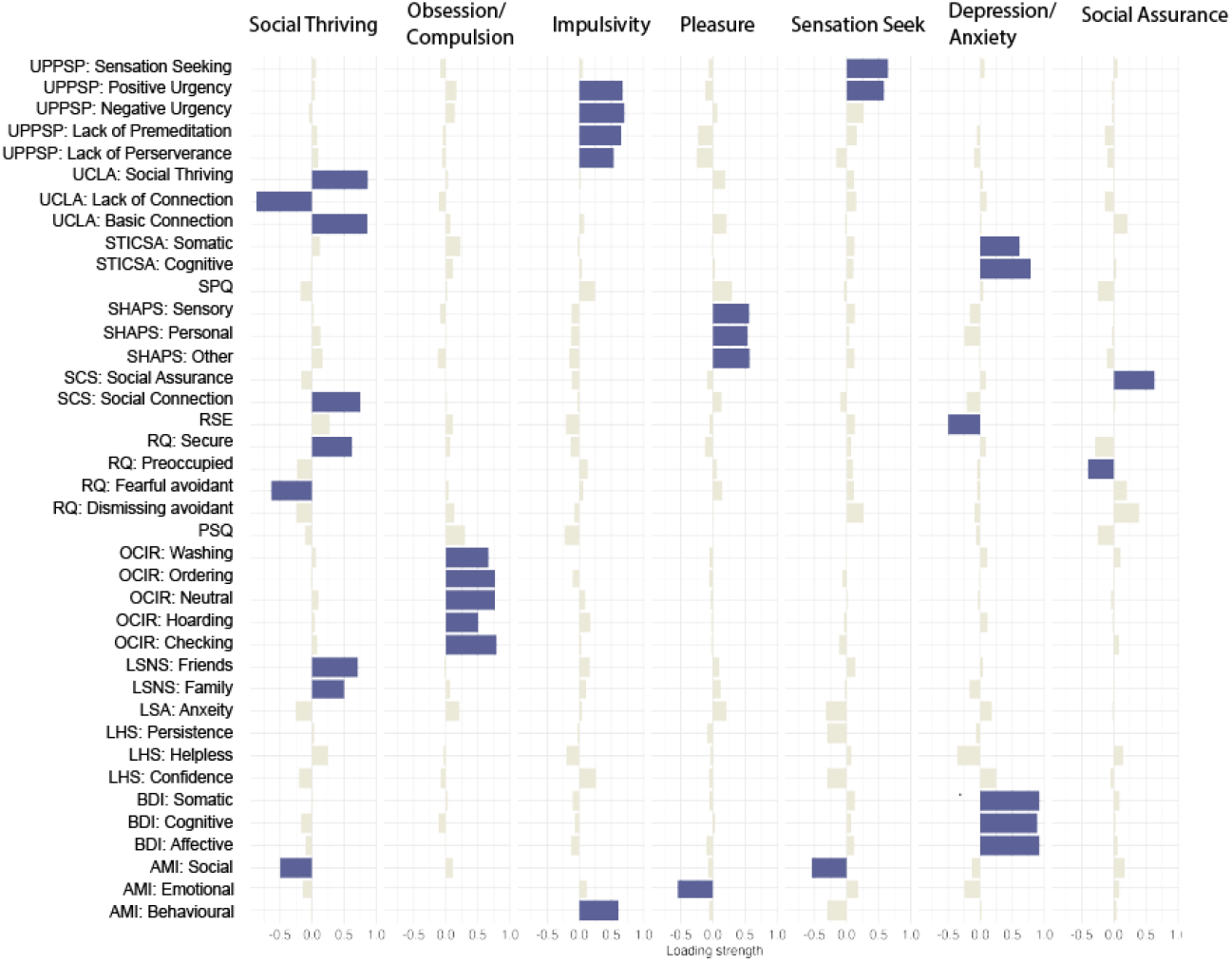
Factor structure and loadings. Results of a factor analysis revealed seven dimensions that captured aspects of participant’s social and non-social personality and psychiatric profile. UPPSP: Urgency, Premeditation (lack of), Perseverance (lack of), Sensation Seeking, Positive Urgency, Impulsive Behaviour Scale. UCLA: UCLA loneliness scale. STICSA: State-Trait Inventory of Cognitive and Somatic Anxiety. SPQ: Social Pain Questionnaire. SHAPS: Snaith-Hamilton Pleasure Score. SCS: Social Connectedness Scale. RSE: Rosenberg Self-Esteem scale. RQ: Relationship Quotient. PSQ: Pain sensitivity questionnaire. OCIR: Obsessive Compulsive Inventory revised. LSNS: Lubben Social Network Scale. LSA: Liebowitz Social Anxiety Scale. LHS: Learned Helplessness Scale. BDI: Beck Depression Inventory. AMI: Apathy Motivation Index.

Based on our preregistration, we focused on three of the seven dimensions extracted from the factor analysis – Social Thriving, Sensation Seeking, and Pleasure (reduced anhedonia). Social Thriving was positively related to scores on the University of California Los Angeles (UCLA) loneliness scale (version 3) ^32^ including the Social Thriving subscale and Basic Connection subscale as well as on the Lubben Social Network Scale’s (LSNS)^33^ Friends subscale and Family subscale, which assess objective social network size. In addition, the social thriving factor we identified had a negative loading on the UCLA Lack of Connection scale and the Apathy Motivation Index (AMI)^34^ Social subscale. Thus, it captured important aspects of people’s social network and connectedness. The Sensation Seeking factor comprised positive loadings on the Sensation Seeking and Positive Urgency subscales of the Urgency, Premeditation (lack of), Perseverance (lack of), Sensation Seeking, Positive Urgency, Impulsive Behaviour Scale^35^ – short (UPPSP) and also a negative loading on the AMI^34^ Social subscale. Thus, it seemed to capture people’s tendency to seek out stimulating positive experiences, possibly in both social and non-social domains. Finally, the Pleasure or reduced anhedonia scale that we identified, had positive loadings for the Snaith-Hamilton Pleasure Score^36^ (SHAPS) Sensory, Personal, and Other-related types of pleasure / anhedonia and a negative loading on the AMI^34^ Emotion subscale. This dimension captured the overall ability to experience pleasure and to seek out pleasant, emotionally positive experiences.

In line with our preregistered hypotheses derived from our discovery data set, we found that participants’ social thriving scores were correlated with the total friend requests they submitted in the task (n=783, r=0.07, p=0.04; Fig.3a). In other words, people who more frequently requested rather than skipped a friendship opportunity were those associated with higher social thriving scores. Participants’ total requests were also related to their sensation seeking trans-diagnostic score (n=783, r=0.13, p=1.25e-4, Fig.3b). We also found that the effect of total requests on social thriving was mediated by sensation seeking (n=783, z=3.1, p=2.1e-3, Fig.3c). A pre-registered hypothesis that social pain was negatively related to total requests did not replicate, because, as noted, the factor itself did not replicate.

**Figure 3.**
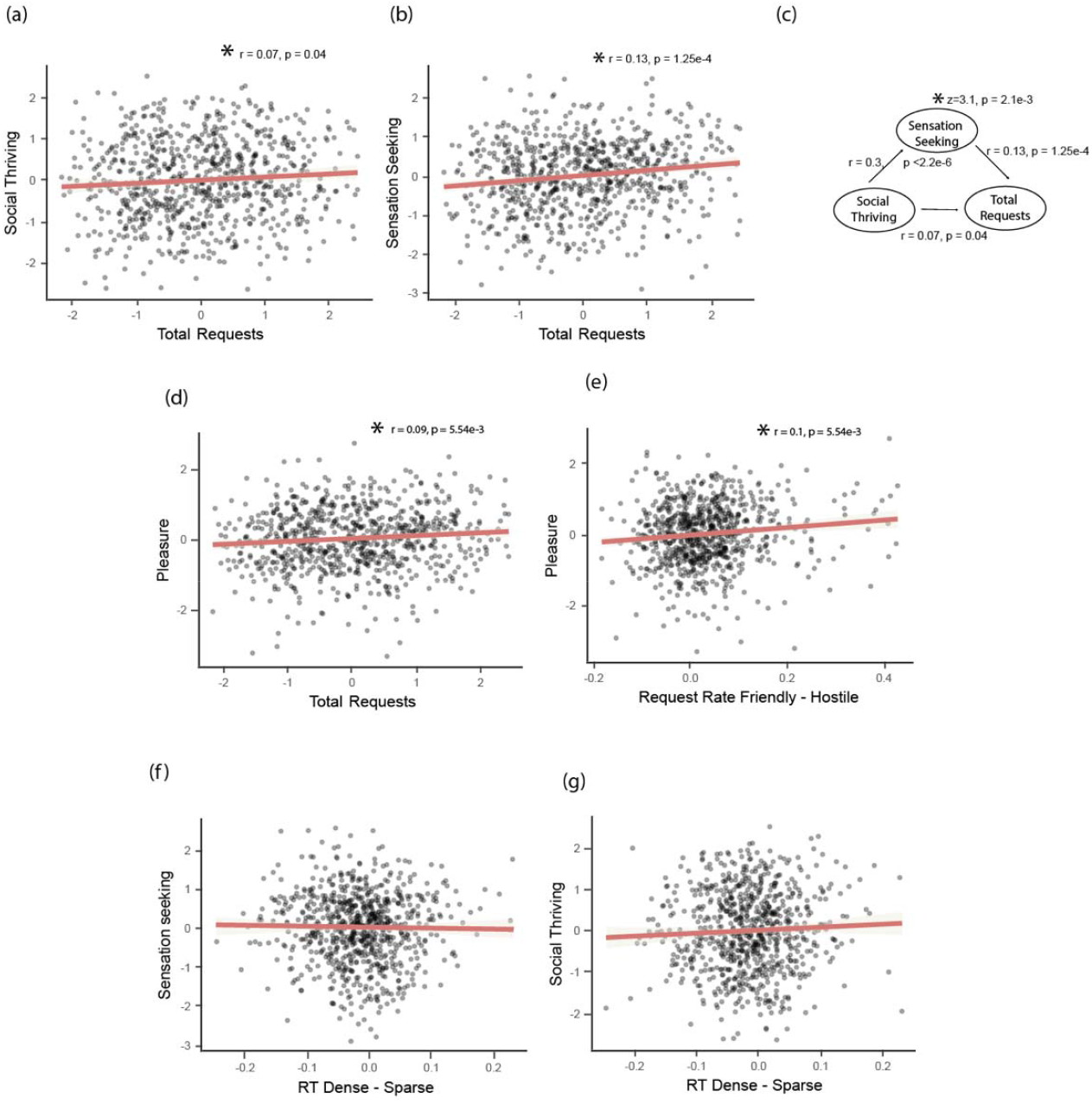
Relationship between social affiliation choices in the Friend Request Task and trans-diagnostic mental health dimensions. (a-b, d-g) Scatter plots show pre-registered relationships between Friend Request Task measures and transdiagnostic dimensions; (c) schematic showing the mediation relationship between social thriving, sensation seeking, and total friendship requests in the Friend Request Task.

Next, the total number of requests made by participants was also related to their pleasure factor score (n=783, r=0.09, p=5.54e-3, Fig.3d). Intriguingly, however, the pleasure factor was not just related to the total number of requests made in the Friend Request Task but, in addition, it was related to the impact of a basic feature of social environment – the average friendliness – on social connection seeking (i.e., the request rate difference between friendly and hostile blocks: n=783, r=0.1, p=5.07e-3, Fig.3e). In summary, of all the dimensions, the Pleasure (or reduced anhedonia) factor, therefore, may be particularly important in capturing how features of the background social environment influence whether people initiate social contact.

While the pre-registered predictions concerning relationships between personality factors and Friend Request Task decision patterns were generally robust, as were RT patterns themselves, the relationships between personality factors and RT patterns were not replicated (Fig3,f-g).

### Statistics of the social environment are tracked by a distributed neural circuit

We employed the same Friend Request Task with a smaller group of 24 participants in a high resolution (1.5mm isotropic), rapid repetition time (1.96 s), accelerated ultra-high field (7T) neuroimaging study. We took care to slightly adjust the task timings (Fig 4a) to allow us to capture the slow blood oxygen level dependent (BOLD) response. We also measured heart rate and respiration during scanning so that we could regress out the effect of physiological noise which is necessary for accurate imaging of the brainstem areas that included some of our regions of interest such as the DRN^13,16,17^ (see Methods). Significant task-related activity patterns in our brain areas of interest, DRN, SN, Hb, aI, and dmPFC, have been identified in similarly sized groups in previous studies using similar ultra-high field imaging protocols^2,15,16^.

**Figure 4.**
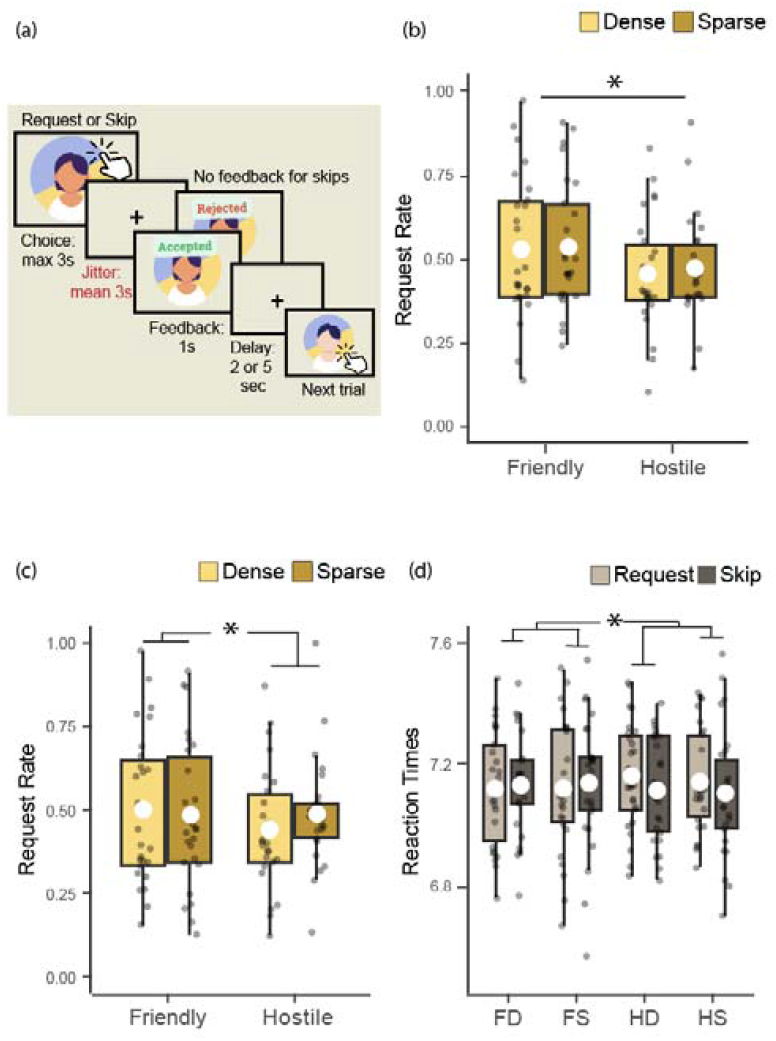
Behavioural results in 7T-fMRI cohort (n=24). (a) Schematic representation of the Friend Request Task with slower timings optimized to capture slow BOLD signals. (b) Boxplots indicate the effect of background social environment on request rates in all trials (white circle indicates mean; box boundaries indicate interquartile range [IQR], encompassing the middle 50% of the data; whiskers extend to furthest data points within 1.5 × IQR from the box boundaries). Request rates were higher in friendly blocks. Y-axis represents request rates, x-axis represents friendliness levels, and colour represents density levels. (c) Effect of background social environments on all but the first 10 trials in each block. Request rates were higher in friendly and sparse blocks. (d) effect of environments on RTs split by action type (request/skip). RTs were faster in friendly blocks, especially when the participant skipped sending a friend request. Y-axis represents RTs, x-axis represents the four types of environments (FD: friendly, dense; FS: friendly, sparse; HD: hostile, dense; HS: hostile, sparse), and colour represents participant action (request/skip).

Before turning to the neural data, first, we checked whether participants in the fMRI experiment exhibited similar behaviour in the Friend Request Task. As before, a 2x2 ANOVA showed a significant effect of friendliness on request rates (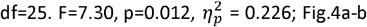), suggesting that people were more likely to send a friend request in friendly environments compared to hostile environments. While there was no effect of density when examining all trials, when we carried out the same check of excluding the first 10 trials in each block that we had performed in the larger sample above, an interaction between friendliness and density was revealed (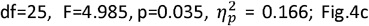). This suggests participants needed an initial period in a task block to appreciate the density of the social environment they were currently in. This seemed particularly important in this slightly slowed-down version of the task.

Looking at the effects of environment statistics on participants’ choices to request or skip sending a friend request, a 2x2x2 ANOVA showed an interaction between friendliness and action on RT (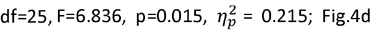). While the main effect of faster RTs in skip trials was not statistically significant in this smaller sample, consistent with the large online dataset, here again, people exhibited a tendency to be faster to skip than to send a request especially in hostile blocks (see supplementary Fig. 2 for results that did replicate).

### Neural encoding of the social environment

Having confirmed that the behavioural results reflected the social environment and largely confirmed our previous results, we next examined where these social factors might be reflected in our *a priori*-defined ROIs. Parameter estimates were extracted from the five ROIs at the time of face onset, separately for all four types of environments. A mixed-model ANOVA showed a main effect of density of the social environment across ROIs (Fig.5a). Sparser blocks were associated with increased BOLD activation at face onset than denser blocks (df =1, *χ*^2^=7.97, p=4.74e-3). There was also a density by region interaction (df=4, *χ*^2^=10.96, p= 2.69e-2). Region-wise post-hoc tests employing a mixed model ANOVA (incorporating factors of both density and friendliness; friendliness effects are discussed below) showed that the interaction was a consequence of a main effect of density in DRN (df=1, *χ*^2^=7.65, p=2.26e-2) and aI (df=1, *χ*^2^=20.50, p=2.98e-5) after correcting for multiple comparisons.

**Figure 5.**
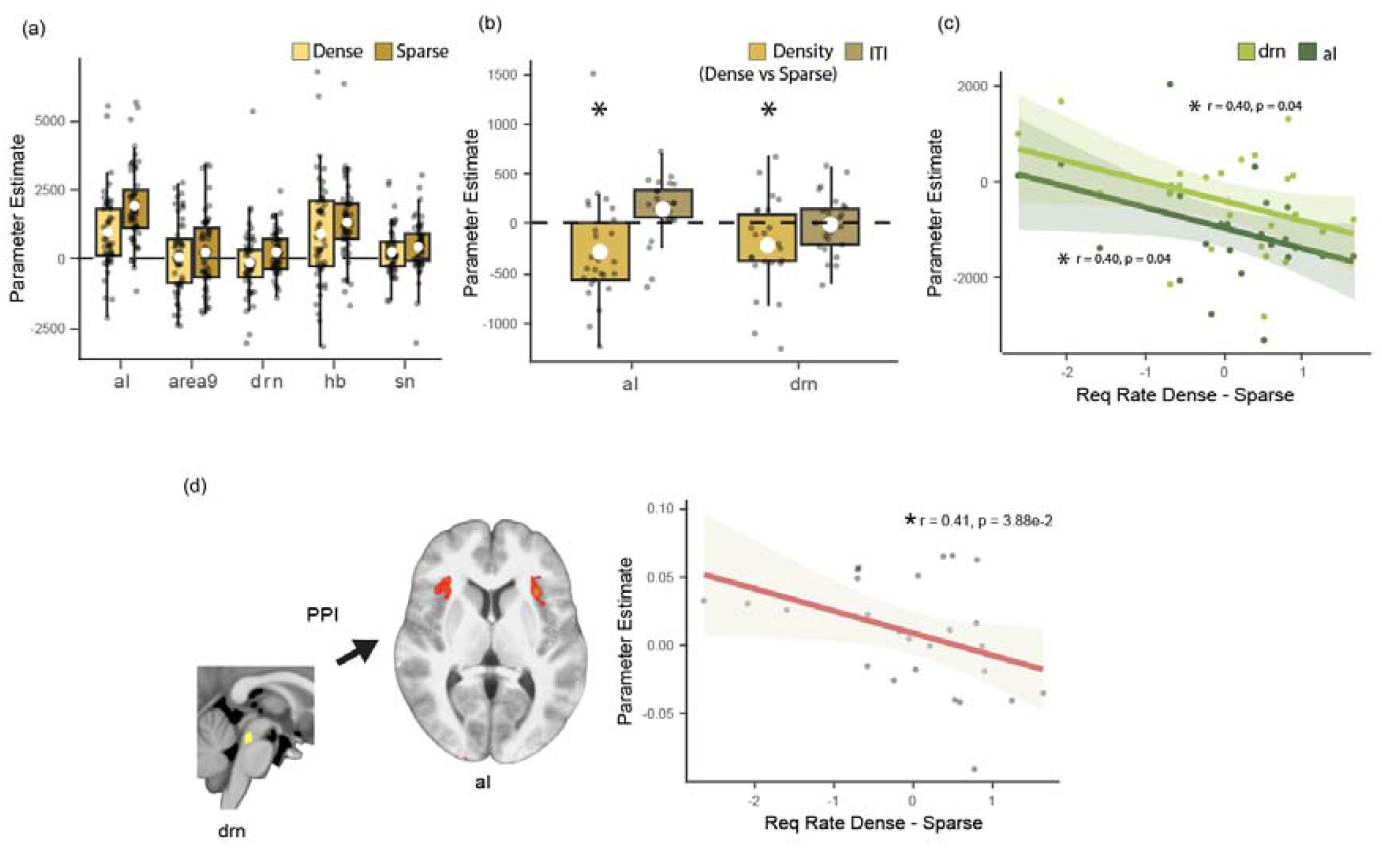
Neural effect of social density. (a) Boxplots indicate the effect of density of social opportunities across five regions of interest (ROIs). X-axis denotes the ROI, y-axis shows parameter estimates, separately for dense (yellow) and sparse (ochre) blocks (white circle indicates mean; box boundaries indicate interquartile range [IQR], encompassing the middle 50% of the data; whiskers extend to furthest data points within 1.5 × IQR from the box boundaries). (b) In the aI and DRN, the two regions where density effects were significant in (a), density effects, shown as one bar reflecting dense vs. sparse (yellow), hold when controlling for local surprise by including past trial ITI (brown). Colour represents explanatory variable from the GLM: yellow= parametric effect of density; brown = previous trial ITI. (c) Across participants, the size of the neural effect of density in aI and DRN relates to the size of the behavioural effect of density. X-axis represents the behavioural effect of density calculated as the difference in request rates observed in dense versus sparse blocks. Colour represents the parameter estimates obtained from DRN (light green) and aI (dark green). (d) Across participants, the PPI effect of density on aI-DRN interactions (PPI effect measured in the aI seeded at the DRN) was correlated with the behavioural effect of density.

Given the density effect on neural activity was mainly driven by the DRN and aI, we further checked whether the effect of density truly reflected the extended experience of long inter-trial intervals (ITIs), and thus the opportunities furnished by the social environment over the duration of the block, rather than simply the length of the most recent interval since encountering the last friend-seeking opportunity. If, instead, neural activity in DRN and aI was simply determined by the most recent ITI then it might be better interpreted in some other way, for example, potentially in relation to the next trial – the next face presentation – starting surprisingly early. We therefore assessed whether an effect of density was still present in these brain regions when controlling for the most recent intertrial interval (ITI), which served as a proxy for local trial-wise surprise. A linear mixed model showed that the density effect remained significant across both regions (df=25, t=-2.19, p=3.84e-2; Fig.5b) even after including the most recent past ITI as additional regressor in the GLM. Further, a mixed model ANOVA showed that the density effect was significantly different from the past trial ITI effect (df=1, *χ*^2^=14.37, p=1.5e-4). Finally, we found that, across participants, the behavioural effect of density was correlated with the neural effect of density both in aI (df=24, t=-2.13, r=0.4, p=4.34e-2; Fig.5c) and DRN (df=24, t=-2.12, r=0.4, p=4.42e-2). We, therefore, finally, examined whether interactions between DRN and aI also varied across participants in a manner that was related to variations in their behaviour. A psychophysiological (PPI) analysis between DRN and aI, which was seeded at the DRN and predicted aI activity as a function of density and DRN activity, showed that the strength of density dependent functional connectivity correlated with the behavioural effect of density on request rates (df=24, r=-0.41, t=-2.18, p = 3.88e-2; Fig.5d).

When we examined the effect of friendliness on neural activity, a mixed model ANOVA identified an interaction between friendliness and action across all areas (df=1, *χ*^2^=8.62, p=3.32e-3; Fig.6ab). The effect of friendliness was stronger in request trials; in other words, BOLD activity differed between friendly and hostile blocks (with greater BOLD responses to face onset in hostile blocks), but it did so more in request trials.

**Figure 6.**
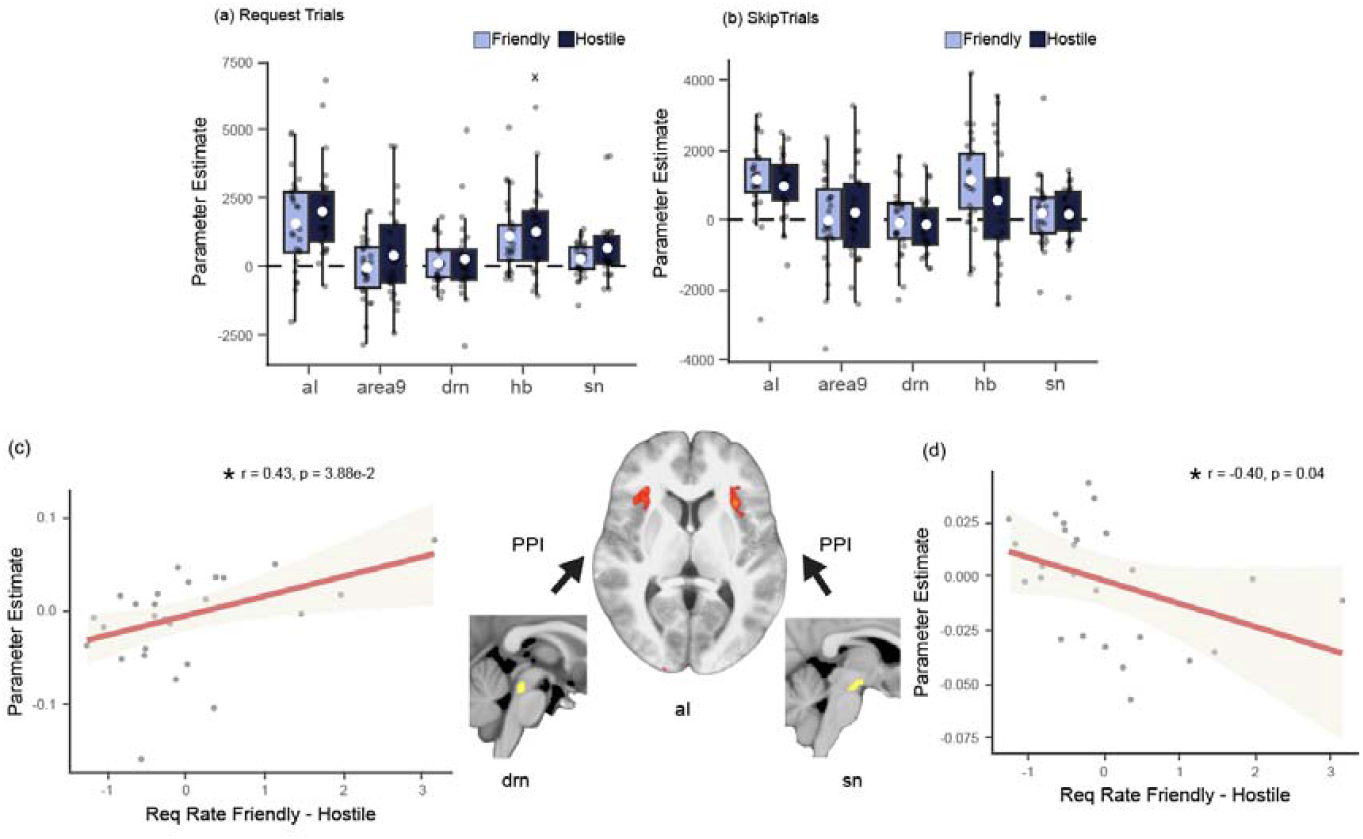
Neural effects of environment friendliness. (a-b) Boxplots indicate the friendliness effects in regions of interest (ROIs) separated by request and skip trials, at the time of face onset. X-axis represents ROI, y-axis represents parameter estimates, and colour represents the levels of friendliness (white circle indicates mean; box boundaries indicate interquartile range [IQR], encompassing the middle 50% of the data; whiskers extend to furthest data points within 1.5 × IQR from the box boundaries). BOLD activity differed between friendly and hostile blocks across ROIS, particularly when sending a friend request; hostile blocks led to greater BOLD responses to face onsets than friendly blocks. (c) Across participants, variation in interactions between DRN and aI as a function of friendliness (PPI effect measured in aI and seeded at the DRN) was related to variation in the behavioural effect of friendliness. (d) Across participants, variation in interactions between SN and aI as a function of friendliness (PPI between SN and aI as a function of the friendliness action interaction effect) was correlated with variation in the behavioural effect of friendliness.

Because we had found a relationship between social environment density and aI-DRN interactions in a PPI analysis, we carried out a similar analysis to assess whether DRN-aI interactions were also linked to the friendliness of a social environment. Again, variation in aI-DRN functional connectivity correlated with variation in the behavioural effect of friendliness on request rates (df=24, r=0.43, t=2.37, p = 2.63e-2; Fig.6c). However, because previous work^16^ had particularly linked one of our other regions of interest, SN, and its interactions with aI, to action initiation, we used a second PPI analysis to examine the relationship between individual variation in SN-aI functional connectivity and individual variation in the impact that friendliness had on action (friend request or skip). SN-aI functional connectivity similarly correlated with the behavioural effect of friendliness on request rates (df=24, r=-0.40, t=-2.15, p = 4.21e-2; Fig.6d).

### SN-aI resting-state connectivity relates to predicted pleasure factor score

So far, we have identified three main groups of findings. First, we found evidence that people’s social connection seeking is influenced by basic statistical features of the social environment such as the density of social encounters and their average friendliness. Second, the same statistics of the social environment were correlated with activity patterns in several brain areas, notably in aI, DRN, and SN and interindividual variations in interactions between aI and the other two areas were associated with interindividual variation in the influence that the statistics of the social environment had on social affiliation seeking. Third, the influence of one statistic of the social environment – mean friendliness – on social affiliation seeking was linked to variation in the Pleasure (or reduced anhedonia) dimension from our factor analysis. In a final investigation we sought to test whether individual variation in this personality variable – Pleasure – was linked to variation in the same neural circuit.

To do this we reasoned that we would need access to a large MRI sample such as that provided by the Human Connectome Project (HCP)^37^. However, we also realized that it was unlikely that such a dataset would include assessment on the same questionnaires that were linked to the Pleasure scale. We therefore examined the questionnaire data that was available in the HCP dataset and attempted to identify any relationship between the measures that were available and our Pleasure factor.

The HCP dataset contains scales measuring related concepts such as the NIH friendship and emotional support toolbox^38^, the Ten item Personality Inventory (TIPI), comprising the Agreeableness, Openness to Experience, Emotional Stability, Extraversion, and Conscientiousness sub-scales^39^. We, therefore, collected scores on the same scales from the 783 participants who performed our behavioural experiments and assessed whether we could use machine learning to predict our factor-derived Pleasure scores from some combination of the NIH friendship and emotional support toolbox and TIPI scores. This was indeed the case (see supplementary Fig. 5), and we were able to create a quantitative translation of the questionnaire scores obtained from HCP participants into predictions of their scores on our Pleasure factor.

We made these behavioural Pleasure factor predictions for n=400 participants in the HCP data set. This subset of HCP n=400 participants was identified based on the availability of both questionnaire scores and cardiac/respiratory recordings that allowed physiological noise clean-up of the BOLD data to achieve more reliable brainstem BOLD signals (see ^40,41^ for more detail). We then examined whether the predicted Pleasure score could be related to variation in resting-state activity between either aI and SN or aI and DRN. We found that our machine learning model-derived Pleasure scores were significantly correlated with the functional connectivity between SN and aI (df=96, r=-0.11, t=-2.34, p=1.97e-2; Fig.7). The greater the connectivity between SN and aI at rest, the lower the participant’s Pleasure score.

## DISCUSSION

Both our own intuitions and a body of research evidence suggest that social connection depends on complex feelings of alignment in peoples’ personal views and perspectives, but also on simple factors such as the individuals’ proximity to and familiarity with one another^4^. Here we demonstrate that, in addition, the fundamental background statistics of social environments impact on participants’ propensities to initiate friendships. Using a novel task design that provided no incentives for or against sending friend requests, we found a significant effect of both the average friendliness of the social environment and the social density of the environment on affiliation request rates. Dense blocks had lower request rates compared to sparse blocks and friendly blocks had higher request rates compared to hostile blocks. The background environments also had a significant effect on RTs. Slower RTs were associated with sparser and friendlier environments. This result was robust across large discovery and replication cohorts and largely replicated in a smaller neuroimaging cohort.

Intriguingly, social affiliation initiation patterns resembled those seen during food-directed foraging^7^, consistent with other recent studies that have observed parallels between social behaviour and behaviours predicted by foraging theories originally constructed in the context of food seeking^42,43^. For example, birds allocate more resources to foraging in abundant environments^7^ which is analogous to people sending more friend requests in friendly environments. Similarly, birds show higher selectivity when the encounter rate with the prey is high^10^ and this is analogous to people sending fewer requests in denser environments.

While drawing analogies between foraging and social affiliation decisions can help us make useful predictions about social behaviour, the parallels with the procedures used in foraging studies are not entirely perfect. For instance, Krebs and colleagues^10^ originally manipulated the value of each choice by offering birds different types of prey (some food items were small while others were large). Our task differs in that all faces carried the same reward value (either a request was accepted or rejected). In the future, we could attribute different values to faces to see how the relative frequencies of different values impact choices^18^.

Similarly, Krebs and colleagues wrapped the food in plastic which the birds had to unwrap before consumption. This served as a clear index for the effort birds had to put in to get their reward. In our study, participants had only to exert a minimal amount of effort to send a friend request (a button press), and consequently there was no clear cost to sending a friend request. While the absence of an effort manipulation makes our task more naturalistic, in the sense that it resembles everyday friend requests on social media, in the future, could the impact of adding an explicit cost for sending requests would be interesting to examine.

In addition to foraging studies conducted in natural environments, our predictions of affiliation seeking were also informed by a review of laboratory-based studies of operant conditioning^20^. This perspective is similar in that it also emphasizes not just the importance of the opportunity that an animal might pursue but also the wider reward environment within which each opportunity occurs. Niv and colleagues point out that by choosing to engage with one opportunity, an animal pays an opportunity cost of not pursuing other opportunities in the background^20^. When the average rate of reward was high, there was a higher opportunity cost to not acting and therefore it was argued that a rat should exert more effort to obtain more reward, and *vice versa* when the reward rate was low.

Such an account, which we refer to as, “the opportunity cost-sensitive model”, might be taken to predict that people should initiate more friendships in a socially dense, as opposed to sparse, environment. It might, however, be difficult to make strong predictions from the opportunity cost-based account given that it is focused more on the speed and vigour of action and on scenarios in which engaging in one opportunity led to decision makers potentially missing out on other opportunities which was not the case in the current study.

While empirical assessments of foraging theory in animal models rarely examine RTs, the opportunity cost-focussed model suggests that environment statistics might directly affect RTs. This is what was indeed the case in the current study. Given that people were disinclined to send friend requests in denser environments, foraging theory predicts they would act more slowly in the interest of being more selective. However, we found that people were faster to act in denser environments in the line with predictions of the opportunity cost-sensitive model.

In our Friend Request Task, we favoured a naturalistic design and chose to employ no explicit cost for sending a friend request. Despite there being no cost for sending a friend request, participants chose to skip sending friend requests about 50% of the time on average. By contrast, some interpretations of rational models of decision making might be taken to predict that, in the absence of costs, people would send a request to every person they encounter, which would in turn maximise both their information gain and their friendships or, in the more limited context of the Friend Request Task, any satisfaction gained from having a friend request accepted. One explanation for the apparently “irrational” behaviour actually observed is that while it is true that there was no explicit, quantified cost for sending a friend request, there was potentially an emotional cost in the form of a rejection which might feel unpleasant. Thus, people might employ an implicit currency of emotional cost when considering making a friend request given that real life experience indicates that there is a limit to the number of social ties we can have and maintain, and the number of rejections we can tolerate^44^. Alternatively, suboptimality in foraging behaviour can be a consequence of stochastic behaviour which, in natural contexts, might nevertheless be useful for acquire information about other courses of action^45,46^. A tendency towards a degree of stochasticity would mean that participants continue to exhibit stochasticity even in the current paradigm despite the absence of information gain.

In this experiment, we defined trans-diagnostic factors to create a simple yet robust profile of participant mental health and personality in a general population sample^47^. Transdiagnostic definitions of mental illness challenge traditional disorder-based frameworks in favour of simple data-driven categories. In general, between-participant variation in the transdiagnostic factors we identified and focused on – Social Thriving, Sensation Seeking, Pleasure (reduced anhedonia) – were related to between-participant variation in one of the Friend Request Task’s simplest parameters – the total requests sent. The relationship between total requests and Social Thriving was mediated by sensation seeking. Such findings suggest both that forming friendships might be part of a larger drive to accumulate sensations or experiences and that variation in these propensities might be captured by a simple laboratory task. Thinking of social affiliation as part of a larger drive is consistent with results suggesting hunger for food and hunger for social contact have shared representations in the brain^6^ and that social animals like macaques trade juice rewards in exchange for the opportunity to gaze at other monkeys^48^.

The transdiagnostic Pleasure factor that we identified was especially intriguing. Not only was it related to the total friend requests that participants made but it was also related to the way in which the social environment impacted on the sending of friend requests; people who had a higher capacity to experience pleasure were also more sensitive to the average friendliness of the social environment and showed stronger adaptation of their friend request tendencies to the environment’s friendliness. In other words, the difference in the number of requests sent in friendly versus hostile environments was more pronounced in these people. Notably, the Pleasure factor included loadings from the apathy motivation index. Given theoretical models linking apathy to the dopaminergic system^20,49^, the pleasure-friendliness link suggested that background friendliness and its interaction with action would be encoded in midbrain dopaminergic areas like the SN. This proved to be the case.

In a 7T fMRI study, we found that background statistics of social environments were tracked by the brain. *A priori* regions of interest (ROIs)—namely the DRN, dmPFC area 9, aI, Hb, SN—encoded opportunities for friendships differently depending on environmental density; that is, at face onset, the activity in these ROIs reflected how often opportunities for affiliation arose. Further, this effect was present even when controlling for the last ITI, ruling out a purely local density code or an interpretation in terms of activity simply reflecting some form of surprise, such as a surprisingly early trial onset. The effect was strongest in DRN and aI. Indeed, a PPI analysis showed that DRN and aI connectivity varied across individuals as a function of the behavioural effect of density suggesting an intimate link between DRN-aI connectivity and mediation of the social density effect. DRN, aI, and DRN-aI connectivity have recently been implicated in tracking the average value of food, monetary rewards, and the potential costs of threats in the environment in macaques and humans^17–19,21^. The present results suggest related roles in tracking the statistics of social environments.

The DRN, while recognized as a serotonergic nucleus, also employs other neurotransmitters. Hence, to test whether the relationship between DRN-aI connectivity and density is specifically linked to serotonin, future studies could manipulate serotonin levels using common drugs such as selective serotonin re-uptake inhibitors.

The brain regions we investigated also encoded an interaction between friendliness and action. A PPI analysis showed that the interaction between friendliness and action (send versus skip friend request) covaried as a function of connectivity between SN and aI. SN and aI-SN interactions have recently been implicated in the integration of opportunity and background value and their translation into action initiation^13,16^. This is consistent with the view that social rewards might be represented in a similar manner in the brain as monetary or gustatory rewards^50^. While the SN is a key source of dopamine, a direct manipulation of dopamine levels, for instance using a dopamine antagonist, might provide insight into the role of dopaminergic neurons in tracking the friendliness of environments.

Finally, we were able to relate individual differences in the aI-SN circuit we had identified with individual differences in the Pleasure scale we had identified. Machine learning was used to make predictions of Pleasure scores for 400 individuals in the HCP data set which were then found to be correlated with aI-SN functional connectivity indices in the same individuals. Such a relationship makes sense given that: 1) the Pleasure factor predicted not just total request rates in the Friend Request Task but the degree to which individuals’ request rates were susceptible to the influence of the average friendliness of an environment; 2) aI and SN activity reflected average friendliness, and aI-SN connectivity predicted variation in the influence average friendliness had on friend requests. Moreover, the findings support the existence of a link between anhedonia, a major symptom associated with depression, and the dopaminergic system^51^. This result is also in line with the role of dopamine neurons in general reward motivated behaviour^52^. It is also consistent with studies showing the involvement of both the dopaminergic system’s^53^ and aI’s^54^ involvement in mediating social approaching and bonding.

## METHODS

### Experiment 1: Large-scale behavioural investigation of friend request seeking and relationships with individual differences in personality and mental health

#### Participants

Ethical approval for this study was obtained through the Medical Sciences Interdivisional Research Ethics Committee (MS-IDREC; ref: R73912/RE001). Informed consent was obtained from each participant before they began the experiment. Multiple datasets were collected as part of this first study, split into a discovery and confirmatory dataset. Participants were recruited using the Prolific recruitment platform (prolific.co). The initial discovery sample included 300 healthy participants out of which 218 participants (mean age = 25.3, males =110, females = 107, other = 1) met the inclusion criteria. For the confirmatory dataset, an a priori power analysis was conducted using G*Power version 3.1^55^ for sample size estimation, based on data from our discovery sample. The smallest size of an effect of interest in the discovery study was 0.017. With a significance criterion of α = .05 and power = .95, the minimum sample size needed would be N = 767 for a t-test computed using the Pearsons correlation coefficient. Thus, the power analysis suggested collecting a minimum sample of size 997 after accounting for a 30% attrition rate. Thus, for the confirmatory dataset, we collected data from 1018 participants out of which 783 met our inclusion criteria (mean age = 27.3, males = 392, females = 380, other = 11). Only participants aged between 18 and 40 years old and who had normal or corrected to normal vision were included in the study.

After collecting pseudonymised data through Prolific, the data were checked for completion. Participants who completed the task were paid at a rate of £6/hr for their participation and they received an additional £2 bonus for task completion. All completed datasets were further examined based on the following preregistered exclusion criteria developed using the discovery sample.

1. Total timeouts (trials in which participants failed to respond after 3 seconds) exceeded 15 in number
2. Request rate per block was either 1 or 0, and that happened for 2 or more blocks out of the total 12.
3. Participants failed to answer repeat questions (a selection of five questions from five different questionnaires presented again towards the end of the experiment) within a 2-point absolute deviation, and this happened for two or more items.
4. Maximum standard deviation of request rates within the same block type was greater than 0.3, suggesting an inconsistent or random choice strategy.
5. Task effects were outside 3 standard deviations of their sample means.

#### Task

A computer-based task was coded using jsPsych^56^ and was uploaded to the Department of Experimental Psychology’s server through “Just Another Tool for Online Studies”^57^ (JATOS) and Pavlovia (pavlovia.org).

In the instructions prior to starting the task, participants were given a cover story in which they were asked to imagine moving to a new city and making new friends. To help them create connections, they would be taken through different clubs which differed in two respects: the friendliness of the people in the club and the number of people that appeared in the club. In each club, they would make repeated choices to send friendship requests to individuals in that club, or to not send a request and skip an opportunity to make a new friend.

Four combinations of clubs were possible: friendly-dense (FD), friendly-sparse (FS), hostile-dense (HD), and hostile-sparse (HS) (Fig.1b). In a friendly club, 80% of all friend requests that a participant sent were accepted whereas in a hostile club, only 20% of all requests were accepted. In a dense club, the time between two consecutive encounters with other club members (being shown a club member face) was briefer; the faces appeared every 2 s with a jitter between 400 and 700 ms. In a sparse club, the time between two consecutive faces was 5 s with an equivalent jitter. On average, a participant saw approximately 32 faces in the dense environments and approximately 19 in the sparse environments.

The background colour of the club (blue or green) indicated the level of friendliness of the club and the background pattern (smaller or larger circles) indicated the level of density of the club. For instance, a blue background with small circles might have indicated a friendly-dense block for some participants. The mapping of backgrounds and patterns to club-type was counterbalanced across participants. At the start of each club, participants were informed about the type of club they would be entering.

Inside each club, participants were shown a series of faces expressing a neutral emotion and given the opportunity to send friend requests. The faces presented as stimuli were obtained from the Chicago Faces Database^58^. The order of the faces was randomised between participants. Faces were sampled at random, and participants saw each face only once.

For every face shown on the screen, participants could either choose to send a friend request by pressing ‘j’ on their keyboard or they could choose to skip by pressing ‘k’. Each face would stay on the screen for as long as the participants took to respond, or for a maximum time of 3 s. If participants sent a friend request, the next screen would show them whether their request was accepted or rejected. This feedback appeared as text over the face and was shown for a duration of 1 s (see Fig.1a for an illustration of the trial structure). If they chose to skip, then a fixation cross of the same duration would appear on the screen. If they failed to respond within the stipulated window, a warning sign appeared on screen indicating they were too slow to respond.

Each block (or club) lasted for a duration of 2.5 minutes. Participants were shown a circular timer on the top right of the screen which indicated the elapsed time. The entire experiment comprised a total of 12 blocks, with all 4 types of blocks appearing three times. The order in which participants encountered these blocks was randomised.

Before the main task began, participants were given a chance to play through one practice block to help them familiarise themselves with the design and controls. After they completed the practice block, their understanding of the task was tested with a short quiz comprising five questions. They were given two attempts to get all questions correct.

At the end of each block, participants were asked three questions. The first question was “How much did you like this club?” and participants indicated their response on a sliding scale from “Not at all” to “Very Much”. Slider position started at the midway point, and participants were required to move the slider at least once before their response could be submitted. The next question was “Did you find a good balance for sending friend requests in this club?” and their responses were taken on a continuous sliding scale from “No, not at all” to “Absolutely yes”. The third and final question was “How do you feel after visiting this club?” and participants indicated their responses on a sliding scale from “Very unhappy” to “very happy”. The purpose of these questions was to get a subjective assessment of how participants viewed the different environments and how they felt while in that club (see supplementary Fig. 1).

During piloting, we noticed that people might not be paying attention to the immediate friendliness of the environment. To make the friendliness salient, we added an attention check. We asked participants “Did the previous person accept or reject your request” roughly every 20 seconds. Participants indicated their responses by pressing ‘f’ on the computer keyboard if accepted, ‘g’ if rejected, and ‘h’ if they skipped the request. This version was used as the task in the discovery dataset. The fact that we observed an effect of friendliness and density of the social environments on acceptance rates suggests that the final task design was sufficiently engaging for participants.

#### Questionnaires

In addition to the behavioural task, participants completed a series of standardised questionnaires to assess their personality and psychiatric profile. Each questionnaire had a lower time limit of 2 s per question to ensure participants took adequate time to understand the questions. If participants completed an entire questionnaire before this lower time limit (2 s x the number of questions), they were given a warning that asked them to reconsider their responses.

We asked participants to complete a set of both social and non-social questionnaires. For obtaining social measures, we used the Lubben Social Network Scale (LSNS) to measure objective social network size^33^. The LSNS comprises two subscales: Friends and Family. Next, we included the UCLA Loneliness Scale Version 3 (UCLA) to measure people’s subjective sense of loneliness^32^. This scale comprises Thriving, Lack of Connection, and Basic Connection as its three subscales. In addition, we included the Social Connectedness Scale (SCS) to measure participants degree of feeling socially connected^59^. The SCS breaks down into Social Assurance and Connectedness subscales. We also used the Relationship Questionnaire (RQ) to measure participants attachment styles^60^. The RQ comprises subscales measuring four attachment styles: Secure, Preoccupied, Fearful Avoidant, Dismissing Avoidant. We also used the Social Pain Questionnaire^61^ (SPQ). Finally, we used the anxiety subscale of the Liebowitz Social Anxiety Scale^62^ (LSA).

Social well-being is also impaired in general mental health conditions like depression and anxiety. To capture such non-social dimensions of well-being, we included the Apathy Motivation Index (AMI) to measure apathy^34^. The AMI comprises Social, Emotional, and Behavioural subscales. In addition, we included Beck’s Depression Inventory^63^ (BDI) and the Snaith-Hamilton Pleasure Score^36^ (SHAPS) to measure depression and its associated symptom of anhedonia. The BDI comprises Somatic, Cognitive, and Affective subscales^64^, whereas the SHAPS comprises Sensory, Personal, and Other-related types of pleasure. We also included the Rosenberg Self-Esteem scale^65^ (RSE) to measure self-esteem and the revised Learned Helplessness Scale^66^ (LHS) to measure learned helplessness. The LHS breaks down into Perseverance, Confidence, and Helplessness subscales.

We included some control questionnaires to assess the specificity of any identified relationships with social or psychiatric dimensions. We acquired data related to symptoms of obsession and compulsion using the checking and neutralising subscales of the Obsessive Compulsive Inventory revised^67^ (OCI). We also recorded impulsivity using the Urgency, Premeditation (lack of), Perseverance (lack of), Sensation Seeking, Positive Urgency, Impulsive Behavior Scale^35^ – short (UPPSP), and anxiety using the State-Trait Inventory of Cognitive and Somatic Anxiety^68^ (STICSA). The UPPSP has Sensation Seeking, Positive Urgency, Negative Urgency, Lack of Premeditation, and Lack of Perseverance as its subscales. Likewise, the STICSA consists of Somatic and Cognitive subscales. Finally, we included the Pain Sensitivity Questionnaire^69^ (PSQ).

Questionnaires were presented in two sets. The first set, comprising SHAPS, UCLA, BDI, LHS, AQ, RQ, PSQ, SPQ, was presented before the behavioural task commenced. The second set, comprising SCS, OCI, LSNS, AMI, UPPSP, LSA, STICSA, was presented after the task ended. In addition, five pre-selected items from these questionnaires were presented again at the end of the study with the intention of using them for data quality checks.

#### Statistical analysis

Data were analysed and figures were plotted using R v4.3.3^70^ running on RStudio v2023.12^71^. Figures were assembled and prepared for publication using Canva (https://www.canva.com) and Adobe Illustrator v28.4.1^72^. All binary categorical variables were coded as -1 and 1. RTs were log transformed. A 2x2 repeated measured analysis of variance (ANOVA) was used to examine the effect of friendliness and density on request rates and RTs. A 2x2x2 repeated ANOVA was used to examine the effects of friendliness and density when RTs were further split by action (requests or skips). A similar 2x2x2 ANOVA was used to examine the effects of past trial feedback, friendliness, and density on request rates. Since all our independent variables only ever had two levels, it was not possible to violate the assumption of sphericity. Significance threshold for all tests were set at p=0.05.

For the factor analysis (Fig. 2), a scree test was used to determine the number of factors to be sought, and the factors were obtained using a promax rotation and minimum residual factoring method. The scores were extracted using the Thurnstone method^73^. A Pearson’s correlation was used to assess the strength of the association between these factors and behavioural measures (Fig. 3). A Sobel test was used to carry out mediation analysis.

Some questionnaires did not come with predefined subscales. For such questionnaires, we undertook factor analysis and defined data-driven subscales. These questionnaires included the SHAPS, which decomposed into sensory, personal, and other-related types of pleasure subscales, UCLA, which decomposed into thriving, lack of connection, and basic connection subscales, and the LHS, which decomposed into perseverance, confidence, and helplessness subscales. Multiple comparisons were not corrected for as the tests were pre-registered.

#### Pre-registered hypothesis

We pre-registered the following hypotheses for main task effects (osf.io/62hw7):

a. People would send friend requests at a higher rate (where request rate is defined as the percentage of trials, out of all trials, on which a request was sent) and will be slower to respond in friendly compared to hostile environments.
b. People would send friend request at a higher rate and will be slower to respond in sparse compared to dense environments.
c. People would send friend requests at a higher rate after acceptance than they will after rejections.
d. People would send friend requests at a higher rate following a previous request particularly in dense environments but will be more likely to make requests following a skip (a non-request) in sparse environments.
e. People would be faster to skip than to request overall. However, this difference between reaction times will be higher in hostile environments than in friendly environments. Similarly, the difference will be higher in dense environments as compared to sparse ones

Distinct factors from personality and mental health questionnaires should be identifiable, that load onto behaviours related to:

- Social Thriving
- Obsession/Compulsion
- Social Pain
- Sensation seeking/Urgency
- Pleasure
- Depression-Anxiety
- Impulsivity

In addition, individual variation in these factors should be predicted by individual variation in task measures. More specifically:

Total requests should relate to factors of social thriving and social pain in the following directions:

- Positively to social thriving
- Negatively to social pain

Total requests should relate to non-social factors in the following manner:

- Positively to sensation seeking
- Positively to pleasure

In a discovery sample, we found that social thriving and sensation seeking were correlated. We then hypothesised that sensation seeking would mediate the relationship between social thriving and total requests. we found a trend (p=0.08) for this mediation using a Sobel test. With a larger sample size, we expected this relationship to hold true. So, we predicted that:

- Sensation seeking would mediate the effect of social thriving on total requests.
- Next, the density effect on reaction times would relate:
- Positively to social thriving
- Positively to sensation seeking

As for the previous hypothesis, we expected the relationship between the density effect on reaction times and social thriving to be mediated by sensation seeking. Finally, the friendliness effect on choices to request or skip social engagement was expected to relate:

- Positively to pleasure

### Experiment 2: 7T high-resolution MRI study of Friend Request Task

#### Participants

The study was approved by the Medical Sciences Division Interdivisional Research Ethics Committee (Ethics Approval Reference: R77443/RE002). Participants were recruited from the Oxford area using the Oxford Participant Recruitment system and other online and email-based advertisement platforms. Prior to attending the scan, participants were screened on the telephone using a standardised Wellcome Centre for Integrative Neuroimaging (WIN) 7T MRI screening form to ensure that they were eligible and safe to be scanned.

We recruited 18–40-year-old right-handed healthy individuals with normal or corrected to normal vision who were fluent in English and 7T-safe. Informed consent was obtained from all participants before the study commenced. Prior to entering the scanner, participants were asked to read through a slideshow which presented instructions about the task and the MRI scanning environment. Participants then changed into scrubs and were screened by a trained radiographer.

A total of 30 participants were recruited out of which 26 (mean age = 24.4, males = 10, females = 16) participants met the inclusion criteria. Behavioural exclusion criteria were similar as those preregistered for the online study (the criteria for the online study were stricter to account for noisy behaviour and questionnaire-related checks). As a result, three participants were excluded because their request rates were either 0 or 1 for two or more blocks. One further participant was excluded because the standard deviation in their request rates exceeded 0.3. The final sample, therefore, comprised 26 participants.

#### Task

While in the scanner, participants played the Friend Request Task described above, albeit with some modifications to make it possible to obtain reliable estimates of brain activity at specific points in each trial given the slow haemodynamic response function (HRF). First, an additional jittered delay (sampled from a gamma distribution centred at 3 s) was added between the response and the outcome to allow dissociation of the HRFs associated with these two events. Next, the block duration was increased from 2.5 minutes to 3 minutes to compensate for the trials that would otherwise have been lost after adding the aforementioned delay (Fig.4a).

Participants indicated their responses by using a button box with two buttons. Participants were instructed to place their index finger over the first button and their middle finger over the second button. The first button was used to send a friend request, whereas the second button was used to skip the opportunity to send a friend request. This button box was placed in the right hand of the participants as they lay inside the scanner. Similarly to the online task, each participant played the task for 12 blocks, with three blocks each of the four different types of environments. The entire task lasted for approximately 45 minutes.

#### MRI protocol

Scans were acquired on a 7T Siemens scanner. Structural scans were obtained using a T1-weighted sequence with a 0.7 mm isometric resolution; GeneRalized Autocalibrating Partial Parallel Acquisition (GRAPPA) acceleration factor of 2; TR 2.2 ms; TE 3.02 ms; 256 slices. The total duration of the structural scan was 6 minutes 35 s. Following the T1 structural scan, field-map images were acquired using a 2 mm isometric resolution; TR 620 ms; TE1 4.08 ms; TE2 5.1 ms; 73 slices; -30-degree angle. The total time to acquire the fieldmap was 2 minutes 2 seconds. Functional scans were then collected while participants performed the Friend Request Task. A multiband sequence was used to obtain a total of 74 slices tilted at a -30-degree angle with a 1.5mm isometric resolution; multiband acceleration factor 2; GRAPPA acceleration factor 2; TR 2.033 s, TE 18.4 ms. The total duration differed slightly between participants depending on their block breaks but was on average 46 minutes 45 seconds. Respiration and heart rate data were acquired while the participants lay in the scanner using instruments manufactured by BIOPAC systems. The total session duration, including set up time, was approximately 75 minutes.

#### Analysis

As for the online behavioural task, a 2 x 2 repeated measured analysis of variance (ANOVA) was used to examine the effect of friendliness and density on request rates and RTs in the behavioural data collected during scanning (Fig. 4b-c). Trials were further split into requests and skips, and a 2x2x2 ANOVA was used to measure the effect of friendliness, density, and action (request versus skip) on RTs (Fig. 4d).

FMRI results were analysed using the FMRIB’s Software Library^74^ (FSL), using the Python programming language^75^ and the package fslpy to interface with FSL^76^. Methods used to plot figures and make statistical inferences were the same as those used in Experiment 1.

Images were prepared by first converting functional runs from dicom file format to nifti and then reorienting them to standard orientation. Structural images were bias-corrected using FSL’s anatomical preprocessing script fsl_anat. Brain extraction was performed using SynthStrip^77^.

Functional images were preprocessed by applying motion correction using FMRIB’s Linear Image Registration Tool^78^ (FLIRT). Fieldmap unwarping was performed using a spatial smoothing parameter set at 3mm of full width half maximum (FWHM). Highpass temporal filtering was also applied. Functional images were registered from their native space to structural space using FMRIB’s Linear Image Registration Tool (FLIRT), and further registered to standard space using FMRIB’s Nonlinear Image Registration Tool^79^ (FNIRT). The Montreal Neurological Institute (MNI) 152 image in 1mm resolution was used as a standard brain template.

After the data was preprocessed, a General Linear Model (GLM) based analysis was run using FMRIB’s Expert Analysis Tool^80^ (FEAT v6). A double gamma haemodynamic response function was used to predict Blood Oxygen Level Dependent (BOLD) responses from task events^81^. The following task based regressors were used in the first GLM (GLM1). For each of the four block types, we used the following three regressors amounting to a total of 12 regressors in the GLM: (1) face onsets, (2) friendliness (parametric), (3) density (parametric), (4) action (request or skip – parametric). All parametric regressors were z-scored. In addition, six basic motion regressors and 33 physiological noise regressors (cosine and sine of basic cardiac and respiratory regressors modelled with an order of 4, and thus 16 regressors; multiplicative cardiac and respiratory terms cos(c + r), sin(c + r), cos(c ™ r), sin(c ™ r), each modelled using an order of 2, and thus again 16 regressors; plus respiration volume per time) were included in the GLM. Physiological noise correction was performed using physiological noise modelling (PNM) tool (https://fsl.fmrib.ox.ac.uk/fsl/fslwiki/PNM), part of the FSL package^82^.

The parameter estimates obtained from fitting the GLM were then extracted for predefined regions of interest (ROI). For all ROIs but the DRN, masks were obtained from similar previous studies^83^ and standard atlases^84^. The DRN mask was drawn using the standard diffusion template included as part of FSL. The rationale for using a diffusion template was the finding that the DRN has lower fractional anisotropy (FA) values and the surrounding tissues have higher FA values, thus leading to an identifiable dark spot (approximate threshold: 3500 where the units are FA*10,000) in diffusion based FA images^85^ (see supplementary Fig. 3; DRN mask also available at https://osf.io/jf6vs/).

A linear mixed-model (LMM) was used to evaluate the effect of friendliness and density on brain activations in five predefined ROIs (Figs.5-6). The LMM included friendliness, density, action, region, and their interactions as fixed effects, and friendliness, density, and action as random effects that varied across participants. The significance of the fixed effects and their interactions was evaluated using a mixed-model ANOVA implementing type II Wald Chi-square tests.

Among the regions that showed the density effect, we tested whether the neural effect was related to the behavioural effect using a Pearson’s correlation. We also tested whether density effects were truly global contextual effects of density, or simply a consequence of the previous ITI as a proxy for local density or surprise. To test for global effects of density above and beyond previous ITI effects, we extended GLM1 to GLM2 which contained identical regressors, but which did not separate them by block-type and included a past trial ITI parametric regressor. Thus, GLM2 included five regressors in total, four regressors analogous to GLM1 (face onset, parametric density, parametric friendliness, parametric action) and a new fifth regressor capturing past trial ITI.

Again, all parametric effects were z-scored. A LMM was used on the parameter estimates to test for the effect of density (against zero; Fig. 5b). Region was included as a fixed effect, and a random intercept was included to account for variability between subjects. A t-test using Satterthwaite’s method was used to assess statistical significance of fixed effects. A mixed model ANOVA was used to determine whether the parameter estimates for density and past trial ITI differed from each other.

Finally, a psychophysiological interaction (PPI) analysis was performed, seeded at the DRN, to test for changes in functional connectivity between DRN and aI as a function of density and friendliness (Fig. 5d,6c). A third GLM was used with face onsets, friendliness, density, action, the DRN time course, and the PPI regressor, which was the interaction between the time course (physiological regressor) of DRN activity and density (psychological regressor). Average parameter estimates were then extracted for the PPI regressor from the aI. A Pearson’s correlation was used to determine whether individual variation in the strength of the PPI estimates was related to individual variation in the behavioural effect of density. An equivalent PPI analysis was performed seeded at the SN to test for functional connectivity with the aI that co-varied as a function of the interaction between friendliness and action (Fig 6d).

### Experiment 3: Large-scale neuroimaging relationships between anhedonia/pleasure and network resting connectivity in the SN-aI network

#### Participants

Two datasets were used in this study. First, for model evaluation and fitting, data from the replication cohort in Experiment 1 were used. Next, for testing the relationship between psychiatric factors and resting-state functional connectivity in the network discovered in Experiment 2, we used the Human Connectome Project (HCP) young-adult database. For this, data and ethics were provided by the WU-Minn Consortium (Principal Investigators: David Van Essen and Kamil Ugurbil; 1U54MH091657; https://www.humanconnectome.org), funded by the 16 NIH Institutes and Centres that support the NIH Blueprint for Neuroscience Research and by the McDonnell Centre for Systems Neuroscience at Washington University. All participants gave informed consent and were reimbursed for their time ($450 for 3T MRI) and travel. HCP participants were scanned at the McDonnell Centre for Systems Neuroscience at Washington University, University of Minnesota (WU-Minn), USA, on a Siemens Skyra 3 Tesla scanner.

For our resting-state analysis, a previously developed resting-state data set from a subset of n=400 HCP participants (males = 207, females = 193, mean age = 29) was used. These participants’ rs-fMRI data had been re-preprocessed by including additional physiological noise regression to improve signal in subcortical regions of relevance here. Motion and ICA components were also regressed out, mimicking the HCP minimal preprocessing pipeline, and all resting-state runs were concatenated. For full details on participant selection and additional preprocessing, see Klein-Flügge and colleagues^40,41^.

#### Task

First, we tried to establish a relationship between HCP participants’ mental health dimensions and those identified in Experiment 1, such as pleasure/anhedonia. To do so, we asked participants from our study in Experiment 1 to complete some of the same questionnaires that participants from the HCP dataset had completed as well. These overlapping questionnaires meant we had some comparable data in both Experiment 1’s large online cohort and the HCP participants. This questionnaire data formed the basis for the first step of our analyses here. We then used machine learning techniques to establish a correspondence between the questionnaire variables we had studied in Experiment 1 and the questionnaire variables obtained in the HCP study.

The questionnaires available in both cohorts that we used were the NIH friendship and emotional support toolboxes^38^ and the Ten item personality inventory (TIPI), comprising the agreeableness, openness to experience, emotional stability, extraversion, and conscientiousness sub-scales^39^. These questionnaires were thought to have the best chance (amongst questionnaires available as part of the HCP dataset) of describing variance related to the social and non-social features that we had found to bear relationships with performance on the Friend Request Task.

#### Analysis

The process of model selection, fitting, and evaluation was performed using the ‘tidymodels’ package in R. First, questionnaire scores were scaled and centred (z-scored) across participants. Then, the dataset was split into training and testing datasets (three quarters of the data were used for training and the remaining quarter was used for testing).

Four types of models were used to predict factor scores from the common set of questionnaires mentioned above: neural networks (nnet), simple linear regression (lm), linear regression with penalised maximum likehood (lmnet), and random forests (rf).

Hyperparameters for the models were tuned using the ‘tuning’ package in R. Models were then compared using the root mean squared error (rsme) and *r*^*2*^ metrics. Winning models selected were those that minimised rsme. Finally, the winning models were validated on the testing dataset using a Pearson’s correlation between the true and predicted values. Model predictions for the Pleasure (or reduced anhedonia) score were obtained for all participants using the winning model.

In our second step, having predicted Pleasure/reduced anhedonia for HCP participants based on their questionnaire data, we turned to their resting-state data. We used all four resting-state runs which had been acquired on a 3T Siemens scanner^86,87^. Each run lasted 14.4 minutes, had a repetition time (TR) of 720ms, echo time (TE) of 33ms, resolution of 2mm isotropic. A total of 72 slices were acquired with a multiband factor of 8 resulting in 1200 timepoints. These data were corrected for distortions, temporally-filtered, minimally smoothed and projected onto a surface reconstruction obtained from aT1-weighted image^87^.

To extract the functional connectivity between SN and anterior insula for each individual, time series were then extracted from the SN and the FOP4^88^ region of the aI using the fully preprocessed concatenated dense rs-fMRI time series of each individual. The aI region used here (FOP4) was chosen as the best match of the insula parcellation provided by Glasser et al.^88^ and the aI region found in our neuroimaging data. A functional connectivity score, indexed by the Pearson’s r between the FOP4 and SN time course, was obtained for all participants to indicate the strength of resting-state connectivity between SN and aI. All further analyses, including model predictions, statistics, and plotting figures, were performed using R v4.3.3^70^ running on RStudio v2023.12. Hypothesised relationships were then evaluated for statistical significance using a simple Pearson’s correlation test (Fig. 7). The significance threshold was set at 0.05.

**Figure 7.**
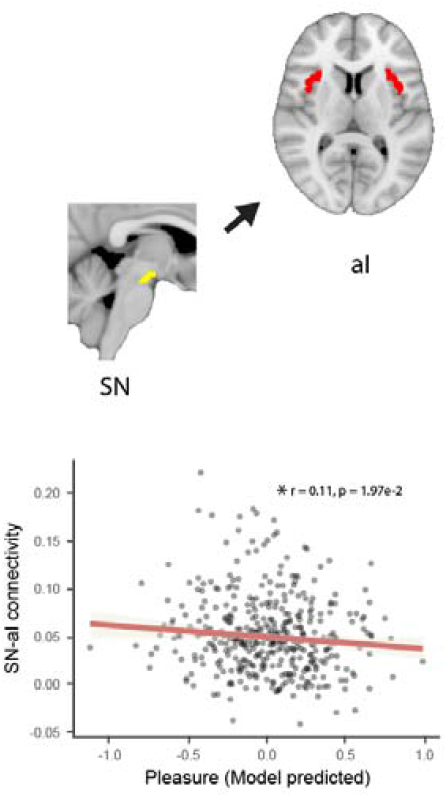
SN-aI resting-state connectivity relates to pleasure (or reduced anhedonia). Functional connectivity between the SN and aI inversely relates to model predicted Pleasure scores in a large group of participants from the Human Connectome Project (HCP; N=400). Greater connectivity between SN and aI was associated with lower pleasure and increased anhedonia.

## ACKNOWLEDGEMENTS

We would like to thank Johannes Algermissen and Simone D’Ambrogio for their help with our study.

## REPORTING SUMMARY

Further information on research design is available in the Nature Research Reporting Summary linked to this article.

## DATA AVAILABILITY

The data reported in the online behavioural experiment as well as the unthresholded statistical group maps of the fMRI experiment are available at the project’s OSF directory (https://osf.io/jf6vs/). All data used in the HCP study are available for download from the Human Connectome Project (www.humanconnectome.org). Users must apply for access and agree to the HCP data use terms (for details see https://www.humanconnectome.org/study/hcp-young-adult/data-use-terms). Here we used both Open Access and Restricted data.

## CODE AVAILABILITY

Code used to analyse behavioural and neural data is available at the project’s OSF directory (https://osf.io/jf6vs/).

## REFERENCES

1. Cacioppo, J. T. & Cacioppo, S. Social Relationships and Health: The Toxic Effects of Perceived Social Isolation: Social Relationships and Health. Soc. Personal. Psychol. Compass 8, 58–72 (2014).

2. Kawachi, I. Social Ties and Mental Health. J. Urban Health Bull. N. Y. Acad. Med. 78, 458–467 (2001).

3. Diener, E. Subjective Well-Being. Psychol. Bull. 95, 542–575 (1984).

4. Hewstone, M. & Stroebe, W. An Introduction to Social Psychology. (John Wiley & Sons, 2021).

5. Baumeister, R. F. & Leary, M. R. The Need to Belong: Desire for Interpersonal Attachments as a Fundamental Human Motivation.

6. Tomova, L. et al. Acute social isolation evokes midbrain craving responses similar to hunger. Nat. Neurosci. 23, 1597–1605 (2020).

7. Pyke, G. H., Pulliam, H. R. & Charnov, E. L. Optimal Foraging: A Selective Review of Theory and Tests. Q. Rev. Biol. 52, 137–154 (1977).

8. Wittmann, M. K. et al. Global reward state affects learning and activity in raphe nucleus and anterior insula in monkeys. Nat. Commun. 11, 3771 (2020).

9. Priestley, L. et al. Dorsal Raphe Nucleus Controls Motivational State Transitions In Monkeys. http://biorxiv.org/lookup/doi/10.1101/2024.02.13.580224 (2024) doi:10.1101/2024.02.13.580224.

10. Krebs, J. R., Erichsen, J. T., Webber, M. I. & Charnov, E. L. Optimal prey selection in the great tit (Parus major). Anim. Behav. 25, 30–38 (1977).

11. Li, S. W., Kietzman, H. W., Taylor, J. R. & Chang, S. W. C. The Evolving Landscape of Social Neuroscience and Its Implications for Psychiatry. Biol. Psychiatry S0006322324013842 (2024) doi:10.1016/j.biopsych.2024.06.004.

12. Gillan, C. M. & Seow, T. X. F. Carving Out New Transdiagnostic Dimensions for Research in Mental Health. Biol. Psychiatry Cogn. Neurosci. Neuroimaging 5, 932–934 (2020).

13. Khalighinejad, N., Priestley, L., Jbabdi, S. & Rushworth, M. F. S. Human decisions about when to act originate within a basal forebrain-nigral circuit. Proc. Natl. Acad. Sci. U. S. A. 117, 11799–11810 (2020).

14. Trier, H. et al. Emotions and individual differences shape foraging under threat. Preprint at 10.31234/osf.io/v6u3y (2023).

15. Khalighinejad, N. et al. A Basal Forebrain-Cingulate Circuit in Macaques Decides It Is Time to Act. Neuron 105, 370-384.e8 (2020).

16. Khalighinejad, N., Garrett, N., Priestley, L., Lockwood, P. & Rushworth, M. F. S. A habenula-insular circuit encodes the willingness to act. Nat. Commun. 12, 6329 (2021).

17. Trier, H. A. et al. A distributed subcortical circuit linked to information-seeking about threat. Proc Natl Acad Sci U A (in press).

18. Khalighinejad, N., Manohar, S., Husain, M. & Rushworth, M. F. S. Complementary roles of serotonergic and cholinergic systems in decisions about when to act. Curr. Biol. CB S0960-9822(22)00104-X (2022) doi:10.1016/j.cub.2022.01.042.

19. Priestley, L. et al. Dorsal Raphe Nucleus Controls Motivational State Transitions in Monkeys. http://biorxiv.org/lookup/doi/10.1101/2024.02.13.580224 (2024) doi:10.1101/2024.02.13.580224.

20. Niv, Y., Daw, N. D., Joel, D. & Dayan, P. Tonic dopamine: opportunity costs and the control of response vigor. Psychopharmacology (Berl.) 191, 507–520 (2007).

21. Wittmann, M. K. et al. Global reward state affects learning and activity in raphe nucleus and anterior insula in monkeys. Nat. Commun. 11, 3771 (2020).

22. Mahmoodi, A., Luo, S., Harbison, C., Piray, P. & Rushworth, M. F. S. Human hippocampus and dorsomedial prefrontal cortex infer and update latent causes during social interaction. Neuron 112, 3796-3809.e9 (2024).

23. Amodio, D. M. & Frith, C. D. Meeting of minds: the medial frontal cortex and social cognition. Nat Rev Neurosci 7, 268–77 (2006).

24. Jamali, M. et al. Single-neuronal predictions of others’ beliefs in humans. Nature 591, 610–614 (2021).

25. Saxe, R. Uniquely human social cognition. Curr Opin Neurobiol 16, 235–9 (2006).

26. Wittmann, M. K., Lockwood, P. L. & Rushworth, M. F. S. Neural Mechanisms of Social Cognition in Primates. Annu Rev Neurosci 41, 99–118 (2018).

27. Wittmann, M. K. et al. Self-Other Mergence in the Frontal Cortex during Cooperation and Competition. Neuron 91, 482–93 (2016).

28. Wittmann, M. K. et al. Causal manipulation of self-other mergence in the dorsomedial prefrontal cortex. Neuron 109, 2353-2361.e11 (2021).

29. Mahmoodi, A. et al. Causal role of a neural system for separating and selecting multidimensional social cognitive information. Neuron 111, 1152-1164.e6 (2023).

30. Mahmoodi, A. et al. A frontopolar-temporal circuit determines the impact of social information in macaque decision making. Neuron S0896-6273(23)00748–1 (2023) doi:10.1016/j.neuron.2023.09.035.

31. Neubert, F. X., Mars, R. B., Sallet, J. & Rushworth, M. F. Connectivity reveals relationship of brain areas for reward-guided learning and decision making in human and monkey frontal cortex. Proc Natl Acad Sci U A (2015) doi:10.1073/pnas.1410767112.

32. Russel, D. UCLA Loneliness Scale (Version 3): Reliability, Validity, and Factor Structure. J. Pers. Assess. 36, ebi (1996).

33. Lubben, J. E. Lubben Social Network Scale. American Psychological Association 10.1037/t10706-000 (2013).

34. Ang, Y.-S., Lockwood, P., Apps, M. A. J., Muhammed, K. & Husain, M. Distinct Subtypes of Apathy Revealed by the Apathy Motivation Index. PLOS ONE 12, e0169938 (2017).

35. Cyders, M. A., Littlefield, A. K., Coffey, S. & Karyadi, K. A. Examination of a short English version of the UPPS-P Impulsive Behavior Scale. Addict. Behav. 39, 1372–1376 (2014).

36. Snaith, R. P. et al. A scale for the assessment of hedonic tone. The Snaith-Hamilton Pleasure Scale. Br. J. Psychiatry 167, 99–103 (1995).

37. Smith, S. M. et al. Resting-state fMRI in the Human Connectome Project. NeuroImage 80, 144–168 (2013).

38. Cyranowski, J. M. et al. Assessing social support, companionship, and distress: National Institute of Health (NIH) Toolbox Adult Social Relationship Scales. Health Psychol. 32, 293–301 (2013).

39. Gosling, S. D., Rentfrow, P. J. & Swann, W. B. A very brief measure of the Big-Five personality domains. J. Res. Personal. 37, 504–528 (2003).

40. Klein-Flügge, M. C. et al. Relationship between nuclei-specific amygdala connectivity and mental health dimensions in humans. Nat. Hum. Behav. 6, 1705–1722 (2022).

41. Jensen, D. E. A., Ebmeier, K. P., Suri, S., Rushworth, M. F. S. & Klein-Flügge, M. C. Nuclei-specific hypothalamus networks predict a dimensional marker of stress in humans. Nat. Commun. 15, 2426 (2024).

42. Gabay, A. S., Pisauro, A., O’Nell, K. C. & Apps, M. A. J. Social environment-based opportunity costs dictate when people leave social interactions. Commun. Psychol. 2, 1–13 (2024).

43. Turrin, C., Fagan, N. A., Dal Monte, O. & Chang, S. W. C. Social resource foraging is guided by the principles of the Marginal Value Theorem. Sci. Rep. 7, 11274 (2017).

44. Dunbar, R. I. M. The Anatomy of Friendship. Trends Cogn. Sci. 22, 32–51 (2018).

45. Trudel, N. et al. Polarity of uncertainty representation during exploration and exploitation in ventromedial prefrontal cortex. Nat. Hum. Behav. 5, 83–98 (2021).

46. Wilson, R. C., Geana, A., White, J. M., Ludvig, E. A. & Cohen, J. D. Humans use directed and random exploration to solve the explore-exploit dilemma. J Exp Psychol Gen 143, 2074–81 (2014).

47. Wise, T., Robinson, O. J. & Gillan, C. M. Identifying Transdiagnostic Mechanisms in Mental Health Using Computational Factor Modeling. Biol. Psychiatry 93, 690–703 (2023).

48. Deaner, R. O., Khera, A. V. & Platt, M. L. Monkeys Pay Per View: Adaptive Valuation of Social Images by Rhesus Macaques. Curr. Biol. 15, 543–548 (2005).

49. Nair, A. et al. Opportunity cost determines free-operant action initiation latency and predicts apathy. Psychol. Med. 1–10 (2021) doi:10.1017/S0033291721003469.

50. Keramati, M. & Gutkin, B. Homeostatic reinforcement learning for integrating reward collection and physiological stability. eLife 3, 1–26 (2014).

51. Belujon, P. & Grace, A. A. Dopamine System Dysregulation in Major Depressive Disorders. Int. J. Neuropsychopharmacol. 20, 1036–1046 (2017).

52. Morales, M. & Margolis, E. B. Ventral tegmental area: cellular heterogeneity, connectivity and behaviour. Nat. Rev. Neurosci. 18, 73–85 (2017).

53. Walum, H. & Young, L. J. The neural mechanisms and circuitry of the pair bond. Nat. Rev. Neurosci. 19, 643–654 (2018).

54. Rogers-Carter, M. M. et al. Insular cortex mediates approach and avoidance responses to social affective stimuli. Nat. Neurosci. 21, 404–414 (2018).

55. Faul, F., Erdfelder, E.Lang, A.-G. & Buchner, A. G*Power 3: A flexible statistical power analysis program for the social, behavioral, and biomedical sciences. Behav. Res. Methods 39, 175–191 (2007).

56. de Leeuw, J. R. jsPsych: A JavaScript library for creating behavioral experiments in a Web browser. Behav. Res. Methods 47, 1–12 (2015).

57. Lange, K., Kühn, S. & Filevich, E. “Just Another Tool for Online Studies” (JATOS): An Easy Solution for Setup and Management of Web Servers Supporting Online Studies. PLOS ONE 10, e0130834 (2015).

58. Ma, D. S., Correll, J. & Wittenbrink, B. The Chicago face database: A free stimulus set of faces and norming data. Behav. Res. Methods 47, 1122–1135 (2015).

59. Lee, R. M. & Robbins, S. B. Measuring Belongingness: The Social Connectedness and the Social Assurance Scales. J. Couns. Psychol. 42, 232–241 (1995).

60. Bartholomew, K. & Horowitz, L. M. Attachment Styles Among Young Adults: A Test of a Four-Category Model. J. Pers. Soc. Psychol. 61, 226–244 (1991).

61. Stangier, U., Schüller, J. & Brähler, E. Development and validation of a new instrument to measure social pain. Sci. Rep. 11, 8283 (2021).

62. Liebowitz, M. R. Liebowitz-social-anxiety-scale - therapist administered.

63. Beck, A. T., Steer, R. A., Ball, R. & Ranieri, W. F. Comparison of Beck Depression Inventories-IA and-II in Psychiatric Outpatients. J. Pers. Assess. 67, 588–597 (1996).

64. Buckley, T. C., Parker, J. D. & Heggie, J. A psychometric evaluation of the BDI-II in treatment-seeking substance abusers. J. Subst. Abuse Treat. 20, 197–204 (2001).

65. Rosenberg, M. Society and the adolescent self-image. Soc. Adolesc. Self-Image 1–326 (1965) doi:10.2307/2575639.

66. Quinless, F. & Nelson, M. Development of a measure of Learned Helplessness. Nurs. Res. 37, (1988).

67. Foa, E. B. et al. The Obsessive-Compulsive Inventory: Development and validation of a short version. Psychol. Assess. 14, 485–496 (2002).

68. Grös, D. F., Antony, M. M., Simms, L. J. & McCabe, R. E. Psychometric properties of the State-Trait Inventory for Cognitive and Somatic Anxiety (STICSA): Comparison to the State-Trait Anxiety Inventory (STAI). Psychological Assessment vol. 19 369–381 (2007).

69. Ruscheweyh, R., Marziniak, M., Stumpenhorst, F., Reinholz, J. & Knecht, S. Pain sensitivity can be assessed by self-rating: Development and validation of the Pain Sensitivity Questionnaire. Pain 146, 65–74 (2009).

70. R Core Team. R: A Language and Environment for Statistical Computing. https://www.R-project.org/ (2024).

71. RStudio Team. RStudio: Integrated Development Environment for R. http://www.rstudio.com/ (2020).

72. Adobe Illustrator. (2024).

73. DiStefano, C., Zhu, M. & Mîndrilã, D. Understanding and Using Factor Scores: Considerations for the Applied Researcher. Pract. Assess. Res. Eval. Pract. Assess. Res. Eval. (2009) doi:10.7275/DA8T-4G52.

74. Jenkinson, M., Beckmann, C. F., Behrens, T. E. J., Woolrich, M. W. & Smith, S. M. FSL. NeuroImage 62, 782–790 (2012).

75. Van Rossum, G. & Drake, F. L. Python 3 Reference Manual. (CreateSpace, Scotts Valley, CA, 2009).

76. McCarthy, P., Cottaar, M., Webster, M., Fitzgibbon, S. & Craig, M. fslpy. Zenodo 10.5281/zenodo.11281355 (2024).

77. Hoopes, A., Mora, J. S., Dalca, A. V., Fischl, B. & Hoffmann, M. SynthStrip: skull-stripping for any brain image. NeuroImage 260, 119474 (2022).

78. Jenkinson, M., Bannister, P., Brady, M. & Smith, S. Improved Optimization for the Robust and Accurate Linear Registration and Motion Correction of Brain Images. NeuroImage 17, 825–841 (2002).

79. Anderson, J., Jenkinson, M. & Smith, S. Non-linear registration, aka spatial normalisation. FMRIB technical report TR07JA2. FMRIB Tech. Rep. TR07JA2 (2010).

80. Woolrich, M. W., Ripley, B. D., Brady, M. & Smith, S. M. Temporal Autocorrelation in Univariate Linear Modeling of FMRI Data. NeuroImage 14, 1370–1386 (2001).

81. Lindquist, M. A., Loh, J. M., Atlas, L. Y. & Wager, T. D. Modeling the Hemodynamic Response Function in fMRI: Efficiency, Bias and Mis-modeling. Neuroimage 45, S187–S198 (2009).

82. Brooks, J. C. W. et al. Physiological noise modelling for spinal functional magnetic resonance imaging studies. NeuroImage 39, 680–692 (2008).

83. Trier, H. A. et al. An ancient subcortical circuit decides when to orient to threat in humans. 2023.10.24.563636 Preprint at 10.1101/2023.10.24.563636 (2023).

84. Pauli, W. M., Nili, A. N. & Tyszka, J. M. A high-resolution probabilistic in vivo atlas of human subcortical brain nuclei. Sci. Data 5, 180063 (2018).

85. Bianciardi, M. et al. Toward an In Vivo Neuroimaging Template of Human Brainstem Nuclei of the Ascending Arousal, Autonomic, and Motor Systems. Brain Connect. 5, 597–607 (2015).

86. Van Essen, D. C. et al. The WU-Minn Human Connectome Project: An overview. NeuroImage 80, 62– 79 (2013).

87. Smith, S. M. et al. Resting-state fMRI in the Human Connectome Project. NeuroImage 80, 144–168 (2013).

88. Glasser, M. F. et al. A multi-modal parcellation of human cerebral cortex. Nature 536, 171–178 (2016).

