## Supplementary Information for "Friend Request Accepted: Fundamental Features of Social Environments Determine Rate of Social Affiliation"

**Additional behavioural results (online and MR cohorts)**

1. Logistic regression mixed-effects model for choices

Here we report all equivalent tests shown in the main part of the manuscript using mixed models.

A logistic regression mixed model ANOVA estimated on the behavioural data of the 783 participants from the online cohort showed similar effects for friendliness and density on choices as reported in the main text. The main effects of friendliness (df=1, $\chi^{2}=82.24,$ p < 2.2e-16) and density (df=1, $\chi^{2}$=164.38, p<2.2e-16) were significant. The interaction between friendliness and density revealed a trend (df=1, $\chi^{2}$=3.09, p=0.08). Further, the interaction between previous action and density was also significant (df=1, $\chi^{2}$=57.46, p=3.45e-14). A full table of the regression results is available in the project’s OSF directory.

2. Mixed-effects models for reaction times

A linear mixed model ANOVA showed the same effects of friendliness and density on reaction times as reported in the main text. The main effects of friendliness (df=1, $\chi^{2}$=54.61, p=1.47e-13) and density (df=1, $\chi^{2}$=59.41, 1.27e-14) were significant, and so was their interaction (df=1, $\chi^{2}$=20.53, p=5.87e-06).

3. Happiness, balance, and liking slider ratings

While participants played the friend request task, we additionally recorded ratings of happiness, balance and liking at the end of each block. The average rating recorded in the online cohort (n=783) are shown in Supplementary Figure 1 below. A 2x2 ANOVA showed a significant effect of friendliness and density on self-reported happiness (friendliness: df=780, F=5485, p<2.2e-16, pes=0.88; density: df=780, F=17.77, p=2.78e-05, pes=0.022; interaction: df=780, F=20.66, p=6.37e-06, pes=0.026), balance (friendliness: df=781, F=3428.15, p<2.2e-12, pes=0.814; density: df=781, F=22.27, p=2.81e-06, pes=0.021; interaction: df=781, F=16.813, p=4.56e-05, pes=0.021), and liking (friendliness: df=782, F=6031.86, p<2.2e-12, pes=0.885; density: df=782, F=9.77, p=2.00e-03, pes=0.012; interaction: df=782, F=31.93, p=2.24e-08, pes=0.039). This shows these two fundamental features of the social environment affected participants in a way that they had awareness over and could report back – adding to the effects on request rates and RTs reported in the main part of the manuscript.

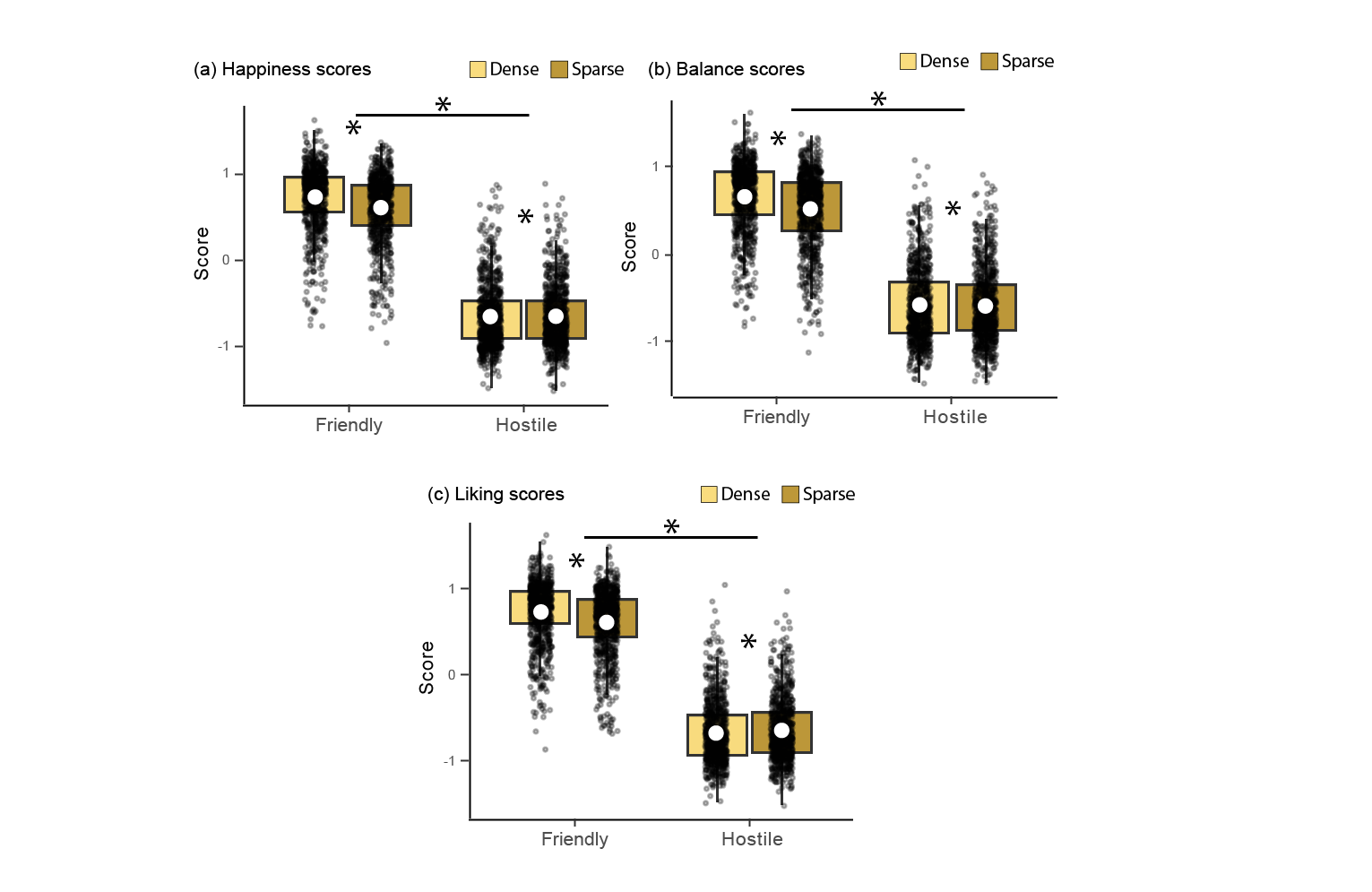

Figure 1. Subjective ratings (z-scored). (a) Happiness, (b) balance, and (c) liking ratings collected at the end of each block (white circle represents the mean, box boundaries indicate the interquartile range [IQR], encompassing the middle 50% of the data; whiskers extend to the furthest data points within 1.5 × IQR from the box boundaries).

4. MRI behavioural effects that did not replicate

While our behavioural results were robust across two large online cohorts, not all results fully replicated in the smaller MR cohort which could be a power issue or related to the slightly slower task timings during the MR data acquisition. For full transparency, the equivalent results significant in the larger cohorts are shown here for the smaller sample.

In the online study, friendliness, density, and interactions had a significant effect on participant reaction time. This effect was not present in the MRI environment (2x2 ANOVA with two levels of friendliness and density; friendliness: df=25, F=0.004, p=0.953, pes=1.44e-4; density: df=25, F=0.462, p=0.503, pes=0.018, interaction: df=25, F=1.10, p=0.304, pes=0.042).

Similarly, in the online study, we found that previous trial outcome had a significant effect on subsequent choices. There was a trend for this effect in the MRI study (2x2x2 ANOVA with two levels of friendliness, density, and outcome; df=24, F=2.94, p=0.099, pes=0.109). In the online dataset, the previous trial outcome also interacted with friendliness of a block. This effect was not significant in the MRI dataset (df=24, F=1.69, p=0.206, pes=0.066).

In a similar vein, previous trial choices (request or skip) also had a significant effect on the request rates, which was not observed in the MRI dataset (2x2x2 ANOVA with 2 levels of friendliness, density, and previous action; df=25, F=0.881, p=0.357, pes=0.034). There was also an interaction between previous trial action and density in the online dataset, which was not observed in the MRI dataset (df=25, F=0.073, p=0.789, pes=0.003).

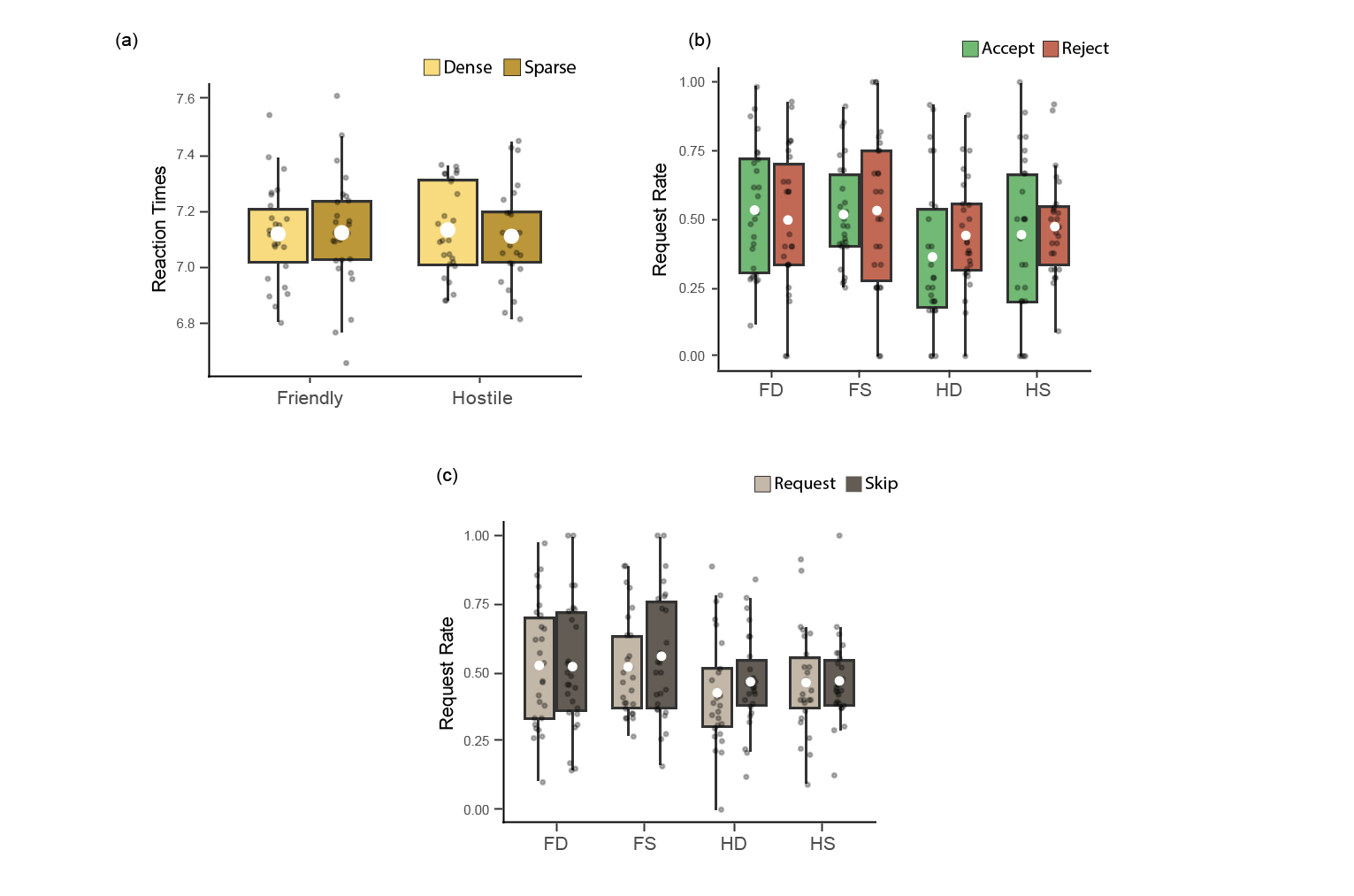

Figure 2: Behavioural effects in the smaller (n=24) MRI cohort for any effects that were significant in the larger online cohort, but not in the MRI study. (a) The effects of friendliness and density on RTs. Y-axis indicates average log-transformed RTs in the respective block type. (b) The effect of previous trial feedback on choice in the subsequent trial in different environment types. Colour indicates feedback received (accept/reject). (c) The effect of previous trial action on choice in the next trial. Colour indicates action (request/skip). Boxplots: white circle within each box represents the mean, while the box boundaries indicate the interquartile range [IQR], encompassing the middle 50% of the data. Whiskers extend to the furthest data points within 1.5 × IQR from the box boundaries.

**Additional results related to the 7T-MRI cohort**

5. DRN mask

We used an anatomically defined mask of the DRN which was drawn on a standard fractional anisotropy (FA) map (FMRIB58_FA) because the DRN is known to have lower FA values than the surrounding tissues, thus leading to an identifiable dark spot (approximate threshold: 3500)^85^ (see supplementary Fig. 3 below).

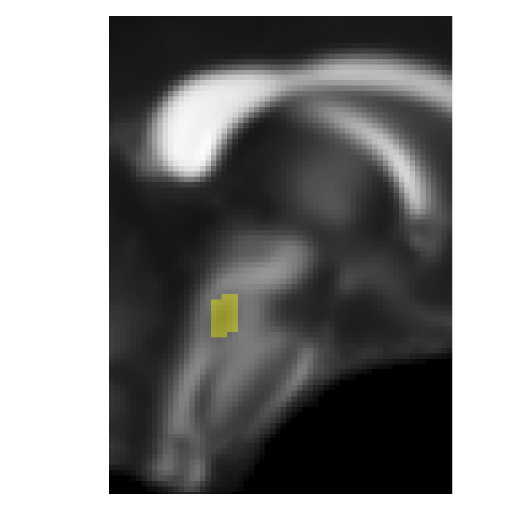

Figure 3: DRN mask overlayed over the standard FSL diffusion template.

6. Friendliness effects at outcome time

In the main part of the manuscript, the effects of the environment’s friendliness were examined at the time of face presentation. Here, we additionally evaluated the effect of friendliness at the time of outcome, given reject/accept decisions may be processed differently as a function of the social environment. Using a mixed model ANOVA, with friendliness, density, region, and their interactions as fixed effects, and friendliness, density, and their interaction as random effects, we found a significant effect of friendliness (df=1, $\chi^{2}$=9.86, p=1.68e-3), and a trend for density (df=1, $\chi^{2}$=2.79, p=0.0947) on parameter estimates encoding outcome. There was also an interaction between friendliness and region (df=4, $\chi^{2}$=27.55, p=1.538e-05). See supplementary Fig. 4 below.

Post-hoc tests were conducted to determine which regions were driving the friendliness effect. They revealed that a main effect of friendliness was present, after correcting for multiple comparisons, in the aI (df=1, $\chi^{2}$=7.60, p=0.02) and area 9 (df=1, $\chi^{2}$=9.36, p=0.01). Thus, we next examined whether the functional connectivity between area 9 and aI was related to model predicted pleasure scores in the HCP dataset (n=400) and found that there was a trend for such an association being present (r=0.1, p=0.05).

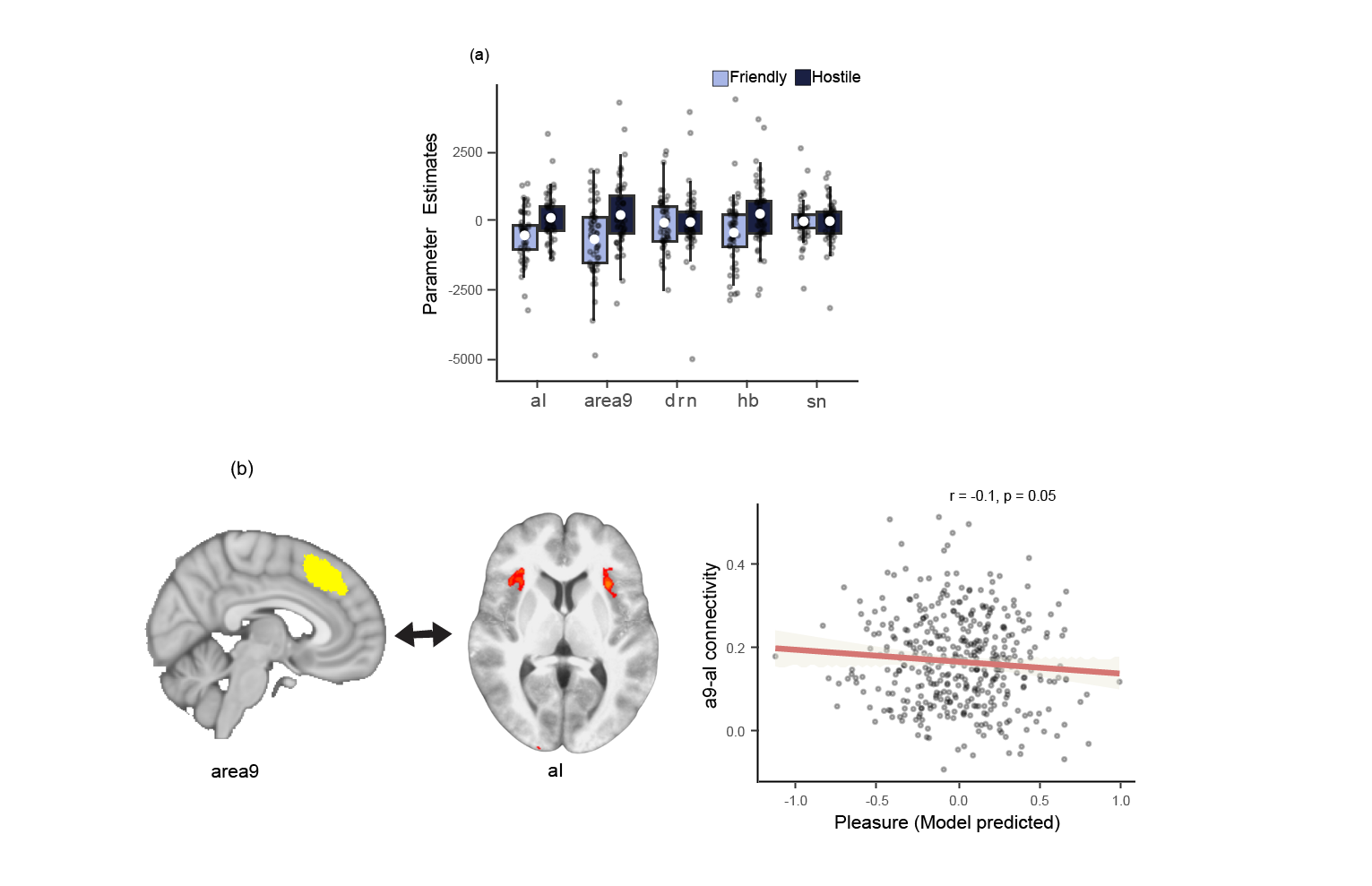

Figure 4: Friendliness effects evaluated at the time of outcome. (a) Boxplots indicate the effect of friendliness on parameter estimates encoding outcome. Colour indicates friendliness. Y-axis represents parameter estimates (white circle within each box represents the mean, while the box boundaries indicate the interquartile range [IQR], encompassing the middle 50% of the data. Whiskers extend to the furthest data points within 1.5 × IQR from the box boundaries). (b) Correlation plot showing model predicted pleasure scores on the x-axis and aI-area9 connectivity on the y-axis.

**Results of the model predictions to derive a Pleasure factor in the HCP cohort**

7. Model evaluations for the pleasure factor

Four models (regularized linear regression, linear regression, neural networks, and random forests) were trained on a training dataset (n=587; obtained from the online Friend Request Task dataset) to predict the pleasure factor score. The winning model was selected that minimised the root mean squared error. The winning model was then evaluated on the testing dataset (n=196; remainder dataset which was not part of the training dataset); the model predicted pleasure factor score was significantly correlated to the true pleasure factor score (r=0.37, p=1.03e-07; see supplementary Fig. 5 below). This predictive model established here was then used to derive a Pleasure-like factor for the HCP participants in the last section of our main manuscript.

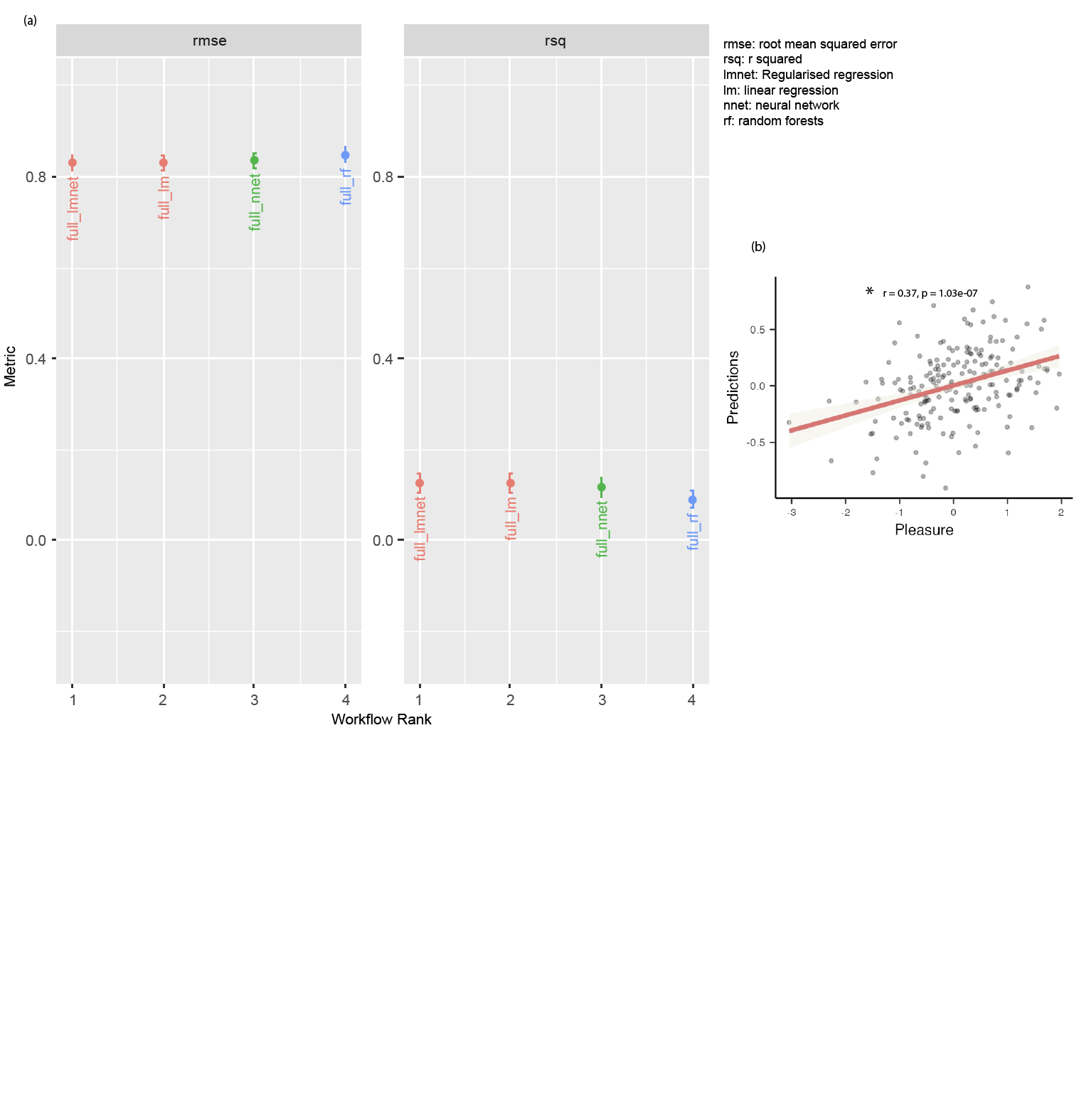

Figure 5: Model evaluations. (a) the left panel shows the root mean squared error, and the right panel shows the r squared for the four models used to predict the pleasure factor in the large online cohort (n=783). (b) correlation between the model predicted pleasure factor (y-axis) and true pleasure factor scores (x-axis), in the test dataset (n=196).
